# Sex-specific DNA methylation signatures of autism spectrum disorder in newborn blood

**DOI:** 10.1101/2024.07.11.603098

**Authors:** Julia S. Mouat, Nickilou Y. Krigbaum, Sophia Hakam, Emily Thrall, George E. Kuodza, Julia Mellis, Dag H. Yasui, Piera M. Cirillo, Yunin Ludena, Rebecca J. Schmidt, Michele A. La Merrill, Irva Hertz-Picciotto, Barbara A. Cohn, Janine M. LaSalle

**Affiliations:** Department of Medical Microbiology and Immunology, School of Medicine, University of California, Davis, CA USA; Perinatal Origins of Disparities Center, University of California, Davis, CA USA; Genome Center, University of California, Davis, CA USA; MIND Institute, University of California, Davis, CA USA; Child Health and Development Studies, Public Health Institute, Berkeley, CA USA; Department of Public Health Sciences, University of California, Davis, CA USA; Department of Environmental Toxicology, University of California, Davis, CA USA; Environmental Health Sciences Center, University of California, Davis, CA USA

**Keywords:** Autism spectrum disorder, DNA methylation, whole genome bisulfite sequencing, newborn blood, newborn dried blood spots, sex differences, female protective effect, epigenetics, X chromosome, differentially methylated regions

## Abstract

**Background:** Autism spectrum disorder **(ASD)** comprises a group of neurodevelopmental conditions currently diagnosed by behavioral assessment in childhood, although neuropathology begins during gestation. A poorly understood male bias for ASD diagnosis is thought to be due to both biological sex differences and cultural biases against female diagnosis of ASD. Identification of molecular biomarkers of ASD likelihood in newborns would provide more objective screening and early intervention. Epigenetic dysregulation has been reported in multiple tissues from newborns who are later diagnosed with ASD, but this is the first study to investigate sex-specific DNA methylation signatures for ASD in newborn blood, an accessible and widely banked tissue.

**Methods:** DNA methylation was assayed from ASD and typically developing **(TD)** newborn blood (discovery set *n* = 196, replication set *n* = 90) using whole genome bisulfite sequencing **(WGBS)**. Sex-stratified differentially methylated regions **(DMRs)** were assessed for replication, comparisons by sex, overlaps with DMRs from other tissues, and enrichment for biological processes and SFARI ASD-risk genes.

**Results:** We found that newborn blood ASD DMRs from both sexes significantly replicated in an independent cohort and were enriched for hypomethylation in ASD compared to TD samples, as well as location in promoters, CpG islands and CpG shores. Comparing females and males, we found that most DMRs with sex differences amongst TD individuals were also found in ASD individuals, plus many additional DMRs with sex differences that were only found in those with ASD. Newborn blood DMRs from females were enriched for the X chromosome and both sexes showed significant overlap with DMRs from umbilical cord blood and placenta but not post-mortem cortex. DMRs from all tissues were enriched for neurodevelopmental processes (females) and SFARI ASD-risk genes (females and males).

**Limitations:** This study is primarily limited by sample sizes, particularly amongst females.

**Conclusions:** Overall, we found a highly replicated sex-specific DNA methylation signature of ASD in newborn blood that showed support for the female protective effect and convergence with epigenetic and genetic signatures of ASD in newborns. These results demonstrate the utility of newborn blood in ASD screening and emphasizes the importance of sex-stratification in future studies.

## Introduction

Autism spectrum disorder **(ASD)** is a group of neurodevelopmental conditions defined by functional impairments in social communication and interactions, combined with elevated restrictive interests and repetitive behaviors (1). Although neuropathology has been shown to begin during gestation (2–7), ASD is typically not diagnosed until early childhood (∼age 3-4) by behavioral observation and clinical assessment (8). Males are diagnosed with ASD approximately four times more frequently than females. This difference may, in part, reflect an underdiagnosis of true cases in females (9) due to diagnostic definitions and tools that primarily consider symptoms more common in males (10–13). For children diagnosed with ASD, intensive behavioral intervention programs have been associated with better outcomes, including higher IQ and improved adaptive behaviors such as communication, daily living skills, socialization, and motor skills (14,15). However, these programs are most effective when begun at an early age, including as young as nine months (16), meaning that children are often diagnosed too late in life to fully benefit. Objective screening tools that use biological measures to identify newborns at increased risk of ASD may help with earlier diagnosis and improved outcomes for many children, particularly females. Insights to biomarker development may be gained by researching newborn tissues and biological markers that have been previously under-studied, such as epigenetic marks from newborn blood.

Several studies have shown that the epigenetic mark of DNA methylation can reflect dysregulation in individuals with ASD, even prior to diagnosis (17–22). DNA methylation is established *in utero*, during which time its landscape is influenced by genetic heritability as well as environmental factors and gene-environment interactions. This triad – genetics, environmental factors, and gene-environment interactions -- also lies at the intersection of etiology for ASD, giving DNA methylation the unique advantage of reflecting multiple early influences of ASD pathology. Intriguingly, ASD studies investigating perinatal tissues such as placenta and umbilical cord blood have shown dysregulated methylation over loci involved in neurodevelopment, demonstrating the promise of perinatal tissues as a surrogate for the brain (17–21). In contrast to cord blood and placenta, newborn blood is accessible and widely banked through newborn dried blood spot **(NDBS)** programs across several states.

Here, we describe the identification of the first sex-specific DNA methylation signatures of ASD from newborn blood using a multi-staged approach **(Figure 1)**. We used NDBS from newborns who were later diagnosed with ASD and sex- and age-matched typical developing **(TD)** children from a California birth cohort (*n* = 196). Following whole-genome bisulfite sequencing **(WGBS)**, which provides the advantage of covering the entire genome, we identified sex-combined and sex-stratified ASD vs TD differentially methylated regions **(DMRs)**. From these DMRs, we identified a sex-specific DNA methylation signature of ASD in newborn blood that was highly replicated in an independent case-control cohort (*n* = 90), showed epigenetic support for the female protective effect, overlapped with ASD DMRs from other tissues (previously published cord blood (17), placenta (21), and cortex (23)), and was enriched for neuro-related biological processes and known ASD-risk genes. Overall, our findings show that newborn blood is an accessible and biologically relevant tissue that can reflect sex-specific methylation signatures of ASD, providing potential insights for biomarker development.

**Figure 1.**
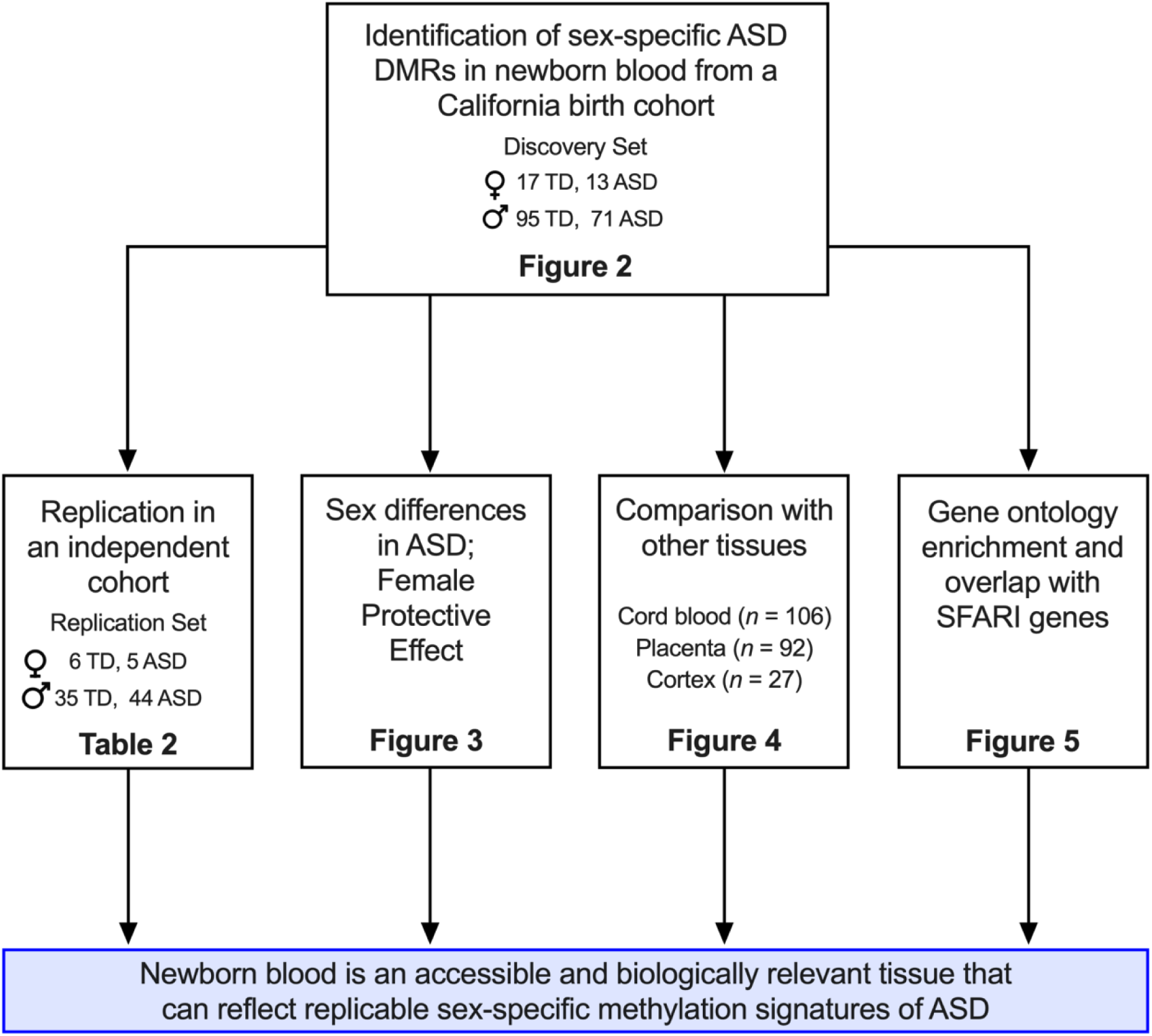
Flowchart of the study.

## Materials and Methods

### Study population

We investigated DNA methylation signatures from ASD and TD individuals using samples from two cohorts of newborn blood (discovery set: 30 females (17 TD, 13 ASD), 166 males (95 TD, 71 ASD); replication set: 11 females (6 TD, 5 ASD), 79 males (35 TD, 44 ASD)), as well as umbilical cord blood (32 females (17 TD, 15 ASD), 74 males (39 TD, 35 ASD)), placenta (30 females (15 TD, 15 ASD), 62 males (31 TD, 31 ASD)), and cortex (10 females (6 TD, 4 ASD), 17 males (5 TD, 12 ASD)). The cord blood (17), placenta (21), and cortex (23) samples have been previously published and are available in the Gene Expression Omnibus with accession numbers GSE140730 (cord blood), GSE178206 (placenta), GSE81541 (cortex) and GSE119981 (cortex). We used the “discovery” data set samples from the cord blood and placenta publications (17,18). Newborn blood, cord blood, and placenta samples were all collected at time of birth, and the post-mortem cortices were collected from individuals aged 4-56 years at the time of death. All samples were collected from different individuals and cohorts, except for placenta and cord blood, which had considerable overlap.

Discovery newborn blood samples were obtained through the Child Health and Development Studies **(CHDS)**, a hospital-based birth cohort that enrolled >20,000 pregnant women at the Kaiser Foundation Health Plan in Oakland, CA from 1959-1967. Over 98% of eligible women enrolled in the study, therefore reflecting the socio-demographics of the Oakland area at the time and making CHDS a more racially diverse cohort than most, with >40% of the selected samples belonging to a racial or ethnic category other than non-Hispanic white (Supplemental Table 1). Follow-up studies have collected information and biological samples from the children and grandchildren of the original cohort. We matched the CHDS database with the California Department of Developmental Services **(DDS)** to identify grandchildren of the original CHDS cohort who have been diagnosed with ASD. TD individuals were chosen from CHDS grandchildren without any DDS diagnoses, matched according to sex and birth year to the ASD individuals. These ASD cases and TD controls were then matched to the California Biobank Program for available NDBS samples, resulting in 196 deidentified NDBS for the discovery group. Replication newborn blood samples were obtained from participants of the CHARGE case-control study in Northern California, which has been previously described (24). For both studies, TD participants are those within the same cohort who did not have any neurodevelopmental disorder listed in the DDS.

Cord blood and placenta samples were obtained through the MARBLES study (25), an enriched-likelihood cohort that enrolls pregnant women in Northern California who already have a biological child with ASD. The TD participants from MARBLES are clinically defined as non-autistic, non-developmentally delayed, having undergone behavioral and cognitive assessments. The human cerebral cortex samples from Brodmann Area 9 were obtained from the National Institute of Child Health and Human Developmental Brain and Tissue Bank for Developmental Disorders at the University of Maryland (23).

### DNA extraction and whole genome bisulfite sequencing

DNA methylation from all samples was assayed using WGBS. Discovery NDBSs were each punched into six 4 mm punches, which were stored at –80C. The samples were randomized by ASD diagnosis and sex and then grouped into ten batches for DNA extraction. Two DNA extractions were performed per sample, each using three 4 mm punches, using the GenTegra Complete DNA Recovery Kit (GenTegra, Pleasanton, CA, USA) and the QIAamp Micro Kit (Qiagen, Hilden, Germany), based on Additional File 2 Protocol GQ from Ghantous et al., 2014 (26). The two isolations from each sample were combined into a single 1.7 μL tube and ethanol precipitation was used to clean the samples. The DNA was re-eluted in 30 μl nuclease-free water and quantified by Qubit. DNA purity was assessed by Nanodrop 260/230 and 260/280 ratios. DNA samples were sonicated to a fragment size of ∼350 bp with a peak power of 175, duty of 10%, and 200 cycles/burst for 47 seconds using 15 μl DNA and 40 μl EB. Sonicated DNA was cleaned and concentrated with gDNA clean and concentrator columns (Zymo Research, Irvine, CA, USA), eluted in 25 μl EB, and re-quantified by Qubit. The maximum mass of DNA for each sample, up to 100 ng, was then bisulfite converted using the Zymo EZ DNA Methylation Lightning Kit.

Illumina sequencing libraries were prepared using the ACCEL-NGS MethylSeq DNA library kit (Swift Biosciences, Ann Arbor, MI, USA), with 6 cycles of indexing PCR (7 cycles for lower input samples). The libraries were pooled and sequenced across 13 lanes of NovaSeq S4 flow cells (Illumina, San Diego, CA, USA) for 150 bp paired end reads with a 5% PhiX spike-in to generate ∼200 million reads (∼8-12x coverage, 60 Gb) per sample. Replication NDBSs were processed and sequenced the same as discovery, except that only one DNA isolation was performed per sample using 2-4 4 mm punches and ethanol precipitation was not used to clean the DNA before library preparation.

Cord blood, placenta, and cortex samples were processed as previously described (17,21,23). Briefly, DNA was extracted from cord blood samples with the Qiagen Puregene Blood kit, and from placental and cortex samples with the Qiagen Puregene kit. DNA from all three tissues was bisulfite converted with the Zymo EZ DNA Methylation Lightning kit. Illumina sequencing libraries were prepared from cord blood and placental samples using the Illumina TruSeq DNA Methylation kit with indexed PCR primers and a 14-cycle PCR program, and from cortex samples as previously described (23). Cord blood and placental samples were sequenced at 2 per lane with 150 base pair paired-end reads and spiked-in PhiX DNA on the Illumina HiSeq X. Cortex samples were sequenced at one per lane with 100 base pair single-end reads on the Illumina HiSeq 2000.

### Descriptive Statistics for the study population

The descriptive statistics compared demographic, parental, and prenatal factors stratified by ASD. This included both categorical and continuous variables, using statistical methods to test for differences between groups. Key variables such as sex, birth year, parental education, race/ethnicity, and insurance status were examined. For continuous variables such as maternal age, a two-sample t-test was used to compare means. For categorical variables such as maternal education, chi-squared tests of independence or fisher’s exact (frequency count < 5) were used to assess associations between the variable of interest and the stratification group. This analysis was performed with R v4.4.1 and SAS v9.4 (Enterprise Edition).

### Sequence alignment and quality control

FASTQ files for each sample were merged across lanes using FASTQ_Me (27) and aligned to the hg38 genome using CpG_Me (28) with the default parameters (29–32). The alignment pipeline includes trimming adapters and correcting for methylation bias, screening for contaminating genomes, aligning to the reference genome, removing PCR duplicates, calculating coverage and insert size, and extracting CpG methylation to generate a cytosine report (CpG count matrix) and a quality control report.

### Determination of Sex

Sex was defined in this study as genotypic sex, whereby those with two X chromosomes were defined as females and those with one X and one Y chromosome were defined as males. Our sample sets did not include any individuals that had other combinations of sex chromosomes. We determined genotypic sex for each sample by calculating the ratio of sex chromosomes from WGBS reads using the SexChecker pipeline (https://github.com/hyeyeon-hwang/SexChecker). The gender identity of study participants is not known to us, and we acknowledge that gender and genotypic sex may not align for some participants. With this manuscript’s goal of better understanding ASD pathogenesis and identifying biomarkers that may be used for newborns, we feel that genotypic sex is an appropriate measure.

### Global methylation analyses

Global methylation for each sample was calculated as the total number of methylated CpG counts divided by the total number of CpG counts from all CpGs included in the DMRichR analysis (filtering described below). Differences in global methylation across tissues were tested with one-way ANOVA with Tukey’s multiple comparisons while differences across sexes and ASD diagnoses were tested with two-way ANOVA with Fisher’s Least Significant Difference using GraphPad Prism 10.0.3.

### Differentially methylated regions

Sex-combined and sex-stratified ASD vs TD DMRs were identified for each tissue (discovery newborn blood, replication newborn blood, cord blood, placenta, cortex), as well as female vs male DMRs for discovery newborn blood using DMRichR (https://github.com/ben-laufer/DMRichR) (33–35). ASD vs TD DMRs included autosomal and sex chromosomes while the female vs male DMRs included only autosomes. We defined regions as sections of the genome that have at minimum 5 CpGs and a maximum gap of 1000 base pairs, with the “universe” for each comparison described as all CpGs included in the analysis. All sex-combined analyses were adjusted for sex, and cortex analyses were additionally corrected for age at time of death. We did not adjust DMRs for cell types due to large inconsistencies across reference datasets (33) and our previous finding that adjustment for cell types did not significantly influence the calling of ASD DMRs in cord blood (17).

Because each tissue and sex contained a different number of samples, we normalized the analyses by adjusting the percent of samples per group that must have at least 1x coverage over a certain CpG site for its inclusion in the analysis. Datasets with fewer samples tend to produce a greater number of DMRs (many of which may be false positives), and datasets with many samples may produce very few (<10) DMRs (which may exclude many real DMRs). To adjust for this, we required all datasets to have the same number of samples covered over a certain CpG for its inclusion in the analysis, which results in a higher percentage of samples for datasets with fewer samples, thereby decreasing the number of DMRs and vice versa for large datasets. Because cortex had the fewest number of samples (*n* = 27), we set perGroup = 1 for all cortex analyses, and for all other tissues, calculated perGroup as 27/*n* so that each comparison required 27 samples to have coverage over a given CpG for its inclusion in the DMR analysis. Otherwise, we used default DMRichR parameters.

### Down sampling males for discovery newborn blood DMR analysis

Because there were a greater number of males than females in the study, discovery newborn blood males were down sampled to match the number of females in order to evaluate the DMRs in each sex when sample sizes were equal. A random number generator was used to select 30 male samples (17 TD, 13 ASD to match the females) for each batch. The “perGroup” value matched for females and down sampled male batches.

### Principal component analysis

DMRs were also visualized using principal component analysis **(PCA)**. Smoothed methylation values over nominally significant (*p* <0.05) DMRs from each comparison were obtained using the “getMeth” function in the bsseq R package v1.36.0. PCA was performed using the “prcomp” function in the stats R package with centering to zero and scaling to unit variance. PCA results were plotted using the “autoplot” function in ggfortify v0.4.16 using R v4.3.1 with an ellipse indicating the 95% confidence level for each group, assuming a multivariate normal distribution.

### Proposed models for sex differences in ASD

The discovery newborn blood dataset was used to investigate three proposed models for sex differences in ASD (36): 1) the multifactorial liability model, including the female protective effect, whereby female-specific protective factors (and/or male-specific vulnerability factors) shift females further away from the liability threshold (37); 2) the extreme male brain theory, whereby all individuals with ASD have a shift towards male phenotypes (38); and 3) the gender incoherence theory, whereby ASD attenuates sex differences present in TD individuals (39).

### Chromosome enrichment analysis

EnrichR (40–42) was used to test discovery and replication newborn blood ASD vs TD DMRs for enrichment across individual chromosomes. The inputs were DMRs mapped to the nearest gene on hg38 with the background set to the “universe” for each DMR comparison, as described above.

### Differentially methylated region overlaps by genomic location and gene name

ASD DMRs were overlapped by genomic location using the Genomic Ranges and RegioneR R packages. The permTest function in the RegioneR package, v1.32.0, was used to calculate significance of overlaps by genomic region as well as mean distance between DMRs of two comparisons. The “genome” was defined as the intersection of all regions included in the two individual analyses. Regions were randomly resampled and tested with 10,000 permutations. For overlap by gene name, ASD DMRs were mapped to the closest gene on the hg38 genome using the “oneClosest” rule of the Genomic Regions Enrichment of Annotations Tool **(GREAT)** (43). Statistics for pairwise comparisons of gene name overlaps were calculated using the Fisher’s Exact Test in the GeneOverlap R package, v1.36.0, in which the “genome” was defined as the intersection of all regions included in the two analyses, mapped to the nearest gene. Numbers of DMR gene name overlaps were visualized using the UpSetR R package v1.4.0 (44).

### Gene ontology term overlaps

Gene ontology **(GO)** enrichment of DMRs from each ASD vs TD comparison were identified with GOfuncR (45), which performs permutation based enrichment testing for the genomic coordinates of the DMRs relative to those of the background regions. Using the sex-stratified DMRs from each tissue, GO terms related to biological processes, molecular functions, and cellular components that had a *p*-value < 0.2 were intersected across all tissues. The overlapping biological processes were graphed by their -log(*p*-value) using GraphPad Prism v 10.0.3, ordered by highest to lowest average -log(*p*-value) across tissue.

### SFARI gene enrichment

DMRs mapped to genes from all comparisons were overlapped with genes listed in the Simons Foundation Autism Research Initiative **(SFARI)** database, release date 3/28/2024 (46). The significance of the overlaps was calculated with the Fisher’s exact test with the background defined as the “universe” for each DMR comparison, as described above.

## Results

### Study subject characteristics

In this study, we investigated WGBS DNA methylation signatures from ASD and TD individuals using newborn blood. We used a discovery cohort of 196 samples [30 females (17 TD, 13 ASD), 166 males (95 TD, 71 ASD)] from the CHDS birth cohort, and replicated our results in an independent cohort of 90 samples [11 females (6 TD, 5 ASD), 79 males (35 TD, 44 ASD)] from the CHARGE case-control cohort. Because ASD is more frequently diagnosed in males and the male:female ratio is decreasing over time, TD controls were matched for both sex and birth year. From the additional information available from birth certificates, only insurance coverage and paternal race/ethnicity were significantly different between ASD and TD within the CHDS cohort, with ASD cases having a higher rate of insurance (91.7% vs. 75.9%, p<0.005), and more Hispanic but fewer non-Hispanic white and non-Hispanic black fathers (p<0.02) than TD (**Supplemental Table 1**). There were no significant differences in the ASD vs. TD samples in the CHARGE replication cohort and both cohorts were demographically diverse **(Supplemental Tables 1-2)**. In both cohorts, global methylation was significantly lower in males than in females but did not differ between ASD and TD samples in either sex (**Supplemental Figure 1**).

### Discovery newborn blood DMRs

We called DMRs for ASD vs TD samples in discovery newborn blood, identifying 59 DMRs in the sex-combined comparison (corrected for sex), 185 in females only, and 99 in males only (Table 1**, Supplemental Figure 2, Supplemental Tables 3-5)**. Upon mapping DMRs to the closest gene on the hg38 genome (54 DMR genes in the sex-combined comparison, 181 in females, 66 in males) and overlapping gene names across the three comparisons, we found that only one DMR gene (*SPATA19*) was present in all comparisons. Of the 181 DMR genes identified only in females, 177 (97.8%) were unique to that comparison. In contrast, 59% (39/66) of DMR genes identified only in the males overlapped with 51.8% (28/54) of those in the sex-combined, primarily due to the large overlap between males only and sex-combined DMR genes **(Figure 2A)**. By genomic location, two DMRs overlapped across females and males, but one of these was methylated in opposite directions in the two sexes, therefore not appearing in the sex-combined analysis. All comparisons showed a signature of hypomethylation in ASD compared to TD samples, but this was more pronounced in the females-only and sex-combined comparisons, where 69% of DMRs were hypomethylated, in contrast to 55% in males **(Figure 2B)**. There was also consistency in the genic and CpG contexts across all comparisons, with significant enrichment for DMRs being in promoters as well as CpG islands and shores in the sex-combined, females-only, and males-only analyses **(Figure 2C-D)**.

**Figure 2.**
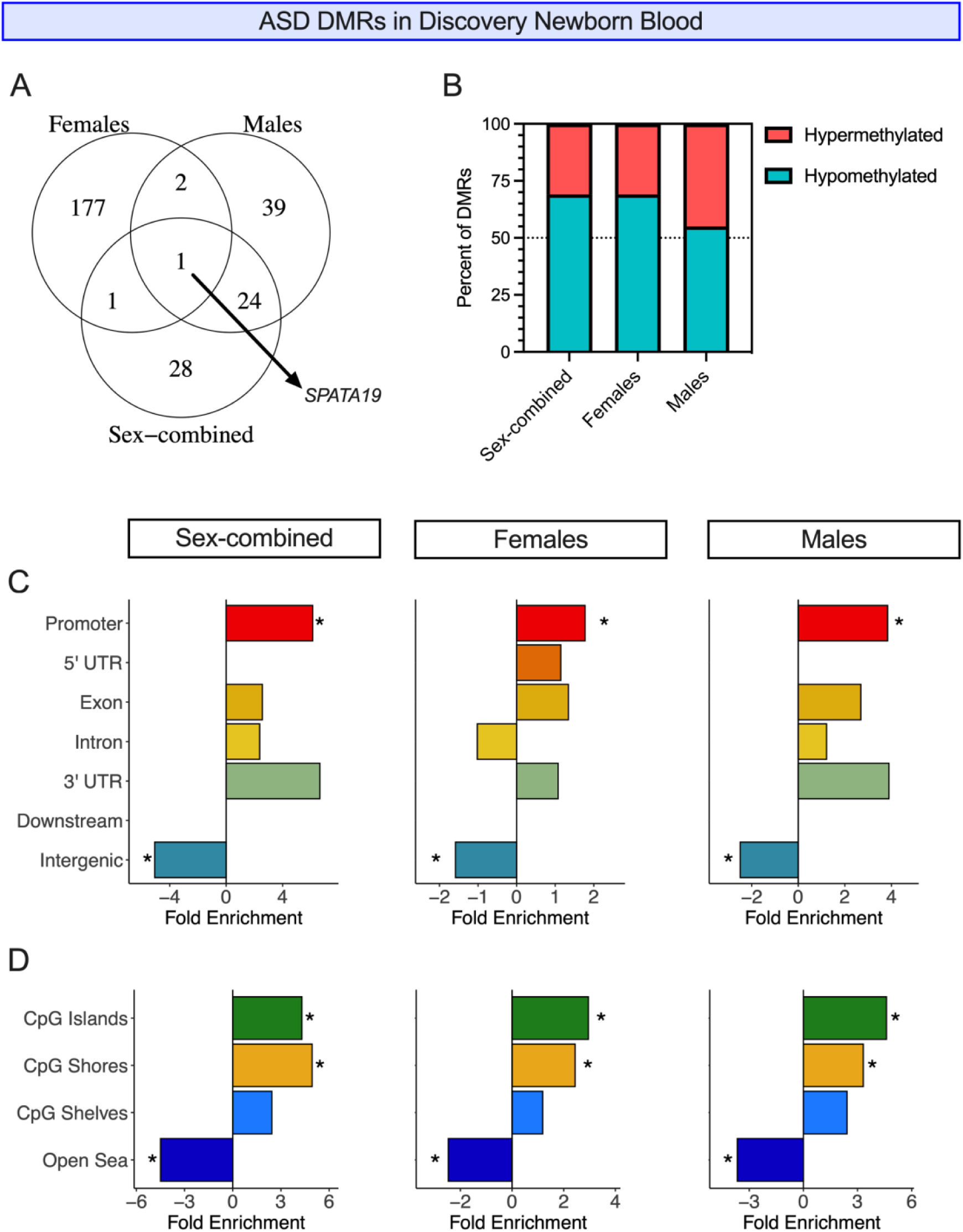
ASD DMRs in discovery newborn blood. **A)** Overlaps between ASD DMRs mapped to the nearest gene on the hg38 genome from sex-combined, females only, and males only comparisons. **B)** The percent of DMRs (permutation *p* <0.05) that are hypo- or hypermethylated in ASD compared to TD samples in sex-combined, females only, and males only comparisons. **C)** Genic and **D)** CpG context enrichments of DMRs (permutation *p* <0.05) from sex-combined, females only, and males only comparisons. DMRs were compared to background regions for each comparison and significance was determined by the Fisher’s test and FDR correction. * = q <0.05.

**Table 1.** ASD vs TD DMRs in discovery newborn blood.

| Comparison | *n* | *n* (ASD) | *n* (TD) | # DMRs | # DMR Genes |
| --- | --- | --- | --- | --- | --- |
| **Discovery Newborn Blood: CHDS birth cohort** |  |  |  |  |  |
| Sex-combined | 196 | 84 | 112 | 59 | 54 |
| Females | 30 | 13 | 17 | 185 | 181 |
| Males | 166 | 71 | 95 | 99 | 66 |
| **Replication Newborn Blood: CHARGE case-control study** |  |  |  |  |  |
| Sex-combined | 90 | 48 | 41 | 67 | 64 |
| Females | 11 | 5 | 6 | 5204 | 4017 |
| Males | 79 | 44 | 35 | 189 | 172 |

Because there were over five times as many males as females in our discovery newborn blood dataset, we also evaluated the epigenetic signature of ASD in subgroups of males that matched the number of samples in females to evaluate the consistency of male DMRs. We randomly subset the males into subgroups of 30 (13 ASD and 17 TD to match the females) and called ASD vs TD DMRs for each batch. We saw that the DMRs separated samples by ASD diagnosis better by hierarchical clustering and PCA in each of the five subgroups of males than when all males were included in a single analysis, matching the females and likely reflecting heterogeneity across male samples **(Supplemental Figure 3)**. Despite this heterogeneity, DMR genes from all five male subgroups significantly overlapped with one another as well as with the DMR genes identified from all 166 males **(Supplemental Figure 4)**, giving us confidence to continue our analysis with the original DMRs identified from all discovery males.

### Newborn blood ASD DMR genes replicated in sex-stratified comparisons

We then evaluated the consistency of epigenetic dysregulation in newborn blood by calling ASD DMRs in an independent cohort, finding 67, 5204, and 189 DMRs in sex-combined, females only, and males only comparisons, respectively **(Table 1, Supplemental Figure 5, Supplemental Tables 6-8)**. When we mapped DMRs to the closest gene on the hg38 genome and overlapped gene lists from discovery and replication analyses, we found significant replication in females-only and males-only comparisons (*p*<0.05) **(Table 2, Supplemental Tables 9-10)**. As a more stringent comparison than gene name overlap, we also compared DMRs from the two cohorts by genomic location. Very few overlaps would be expected by chance since DMRs cover a very small fraction of the genome and indeed, there were fewer overlaps than by gene name in all comparisons (1 in sex-combined, 12 in females only, and 4 in males only). However, all were significant (permutation *p*-value < 0.05) **(Supplemental Tables 9-10REF)**. We also analyzed whether, given a DMR in the discovery group and its closest DMR by location in the replication group, the DMRs were closer together in the genome than would be expected by random chance and found that this was true for all comparisons (permutation *p*-value=0.0001) **(Supplemental Figure 6**, **Supplemental Table 9)**.

**Table 2.** Replication of newborn blood ASD vs TD DMR genes.

| Comparison | Discovery Set: Number of DMR Genes | Replication Set: Number of DMR Genes | Overlap | Percent Overlap | Jaccard Index | Odds Ratio | $p$ value |
| --- | --- | --- | --- | --- | --- | --- | --- |
| Sex-combined | 54 | 64 | 1 | 1.85% | 0.0085 | 4.55 | 0.203 |
| Females | 181 | 4017 | 87 | 48.07% | 0.021 | 2.58 | **4.79E-10** |
| Males | 66 | 172 | 7 | 10.61% | 0.030 | 10.97 | **8.26E-06** |

### Discovery newborn blood DMRs show support for the female protective effect

Because only three ASD DMR genes overlapped between females and males, we wanted to further explore potential sex differences in our discovery newborn blood dataset. The skew in ASD diagnoses towards males has led to several theoretical models for neurobiological sex differences in ASD (36), including, 1) the multifactorial liability model, including the female protective effect (37); 2) the extreme male brain theory (38) and; 3) the gender incoherence theory (39) **(Figure 3A)**. We evaluated whether there was epigenetic support for these theories in newborn blood.

**Figure 3.**
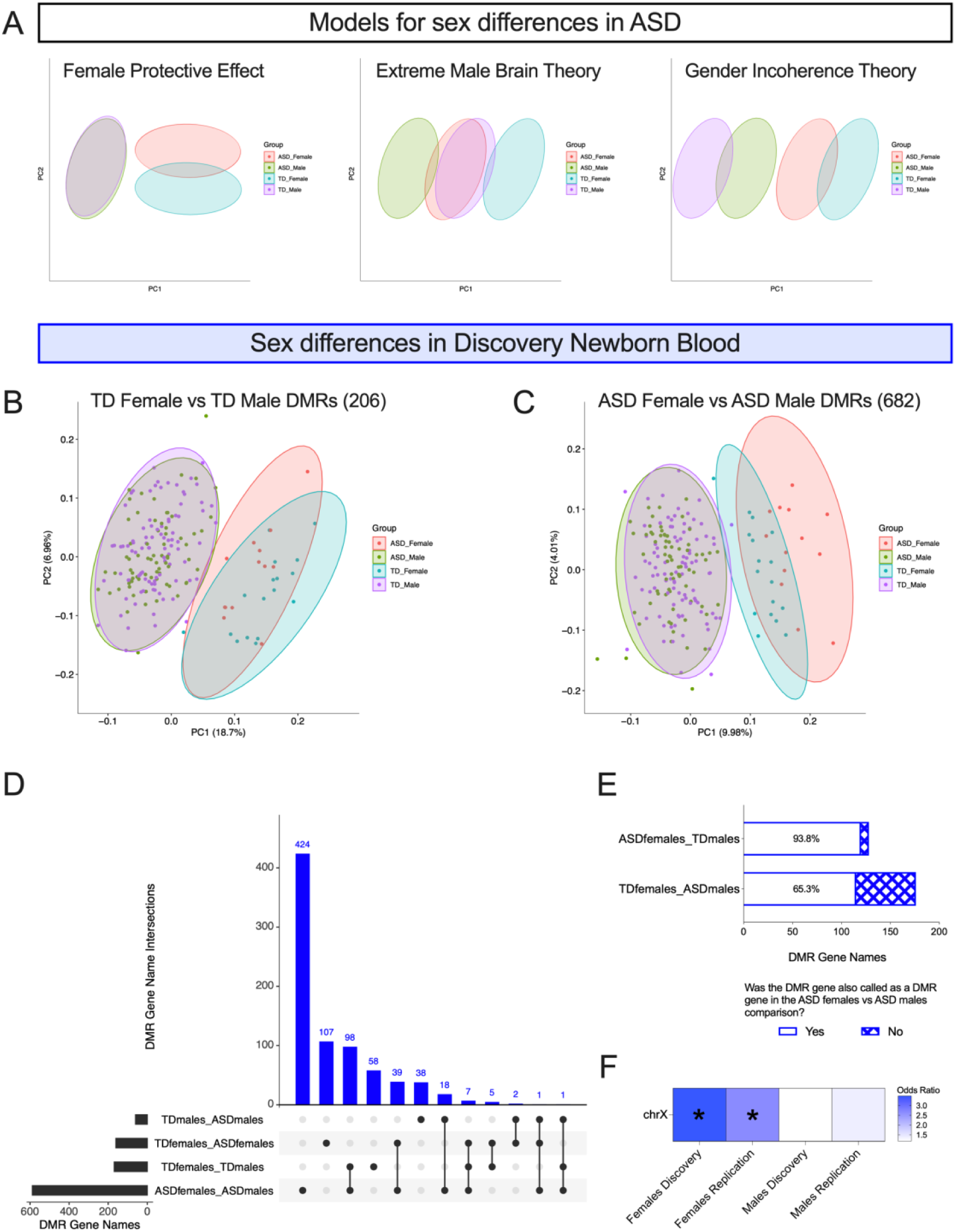
Discovery newborn blood DMRs display female protective effect. **A)** PCA models reflecting different theories for sex differences in ASD. PCA of smooth methylation values in discovery newborn blood samples over DMRs from **B)** TD female vs TD male comparison and **C)** ASD female vs ASD male comparison. **D)** UpSet Plot of autosomal DMRs intersected by gene name. **E)** Bar plot of ASD female vs TD male and TD female vs ASD male comparisons, showing proportion of autosomal DMR genes that were also called as an autosomal DMR gene in the ASD female vs ASD male comparison. **F)** Heatmap showing enrichment on the X chromosome from ASD vs TD DMRs identified from discovery and replication newborn blood females and males. \**q* <0.05

We first identified autosomal DMRs between females and males within TD (TD females vs TD males) **(Table 3)** and within ASD (ASD females vs ASD males) **(Supplemental Table 11-12)**. To understand how sex differences intersect with the methylation signature of ASD, we performed PCA from smoothed methylation values from all discovery newborn blood samples over the diagnosis-specific female vs male DMRs and found that ASD males did not separate from TD males in either comparison, while ASD females showed some separation from TD females, though the direction of separation depended on the regions assayed: when smooth methylation was assayed over TD female vs TD male DMRs **(Figure 3B)**, the ASD females were shifted towards the males, while the opposite occurred when methylation was assayed over ASD female vs ASD male DMRs **(Figure 3C)**.

**Table 3.**
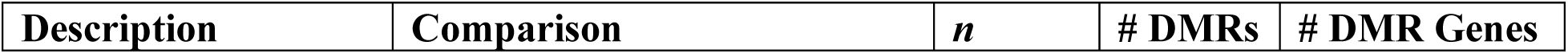
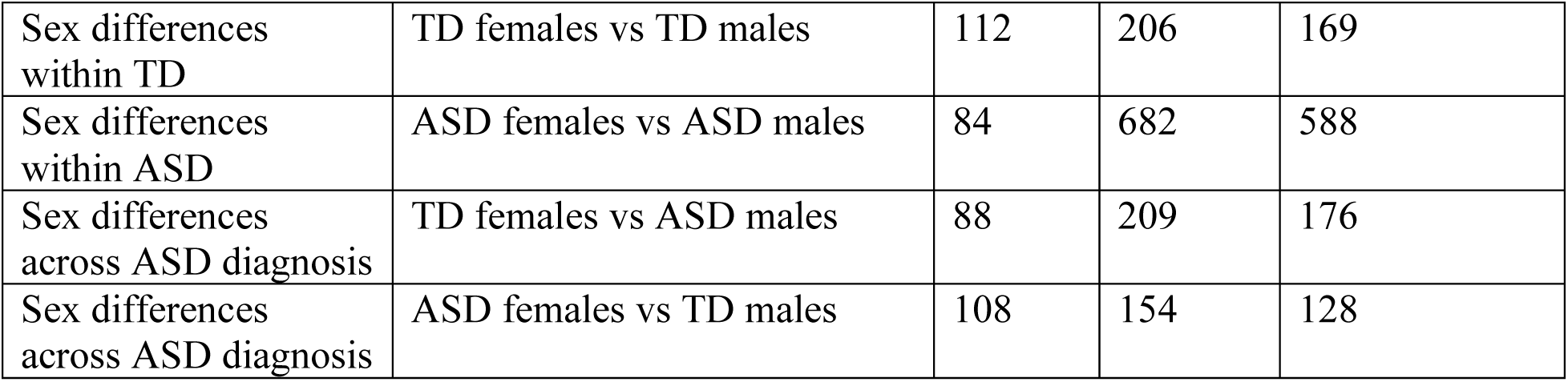
Female vs male DMRs in discovery newborn blood autosomes.

| Description | Comparison | *n* | # DMRs | # DMR Genes |
| --- | --- | --- | --- | --- |
| Sex differences within TD | TD females vs TD males | 112 | 206 | 169 |
| Sex differences within ASD | ASD females vs ASD males | 84 | 682 | 588 |
| Sex differences across ASD diagnosis | TD females vs ASD males | 88 | 209 | 176 |
| Sex differences across ASD diagnosis | ASD females vs TD males | 108 | 154 | 128 |

We next overlapped autosomal DMR gene names from ASD vs TD and female vs male comparisons to better understand the proportion of methylation differences that are common versus unique across sexes and diagnoses **(Supplemental Table 13)**. We found that the majority (62.7%; *p* = 2.3E-106) of TD female vs TD male DMR genes were also found in the ASD female vs ASD male comparison, with many additional DMRs in the latter comparison **(Figure 3D) (Supplemental Figure 7)**. In addition, of the 128 ASD female vs TD male DMR genes, 120 (93.8%) were also called as ASD female vs ASD male DMR genes, indicating that epigenetic sex differences are consistent regardless of ASD diagnosis in the males **(Figure 3E)**. This was not true when examining the epigenetic impact of ASD diagnosis in females, as only 115/176 (65.3%) of the TD female vs ASD male comparison DMR genes were also called as ASD female vs ASD male DMR genes. These results support our previous finding across perinatal tissues that the signature of ASD is more distinct in females than in males, a primary prediction of the multifactorial liability threshold and female protective effect model (36).

### Discovery and replication newborn blood ASD DMRs identified from females are enriched for X chromosome location

Because we saw evidence of the female protective effect from our analysis of autosomal chromosomes, we wondered if methylation differences on the X chromosome were further contributing to the signature of ASD in females. Using both the discovery and replication newborn blood datasets, we tested the ASD vs TD DMRs identified from females only, males only, and sex-combined comparisons for enrichment across all chromosomes. We found that DMRs identified from females in both cohorts were significantly enriched for X chromosome location (*q* <0.05), while DMRs identified from males only or sex-combined analyses were not significantly enriched for any chromosome **(Figure 3F) (Supplemental Table 14)**. Consistent with this finding, none of the DMRs derived from the down-sampled male subgroups from the discovery set were enriched for any chromosome. Additionally, of the twelve loci that replicated in females across both newborn blood datasets, three (25%) were located on the X chromosome **(Supplemental Table 10)**. These findings indicate that the methylation signature of ASD in females is being driven by both autosomal and X-linked loci.

### Newborn blood ASD DMR genes significantly overlap with DMR genes from umbilical cord blood and placenta

Once we established the sex-specificity of the ASD epigenetic signature in newborn blood, we tested if the ASD vs TD DMRs overlapped with those identified from other tissues. We called DMRs from previously published umbilical cord blood, placenta, and post-mortem cortex WGBS data from ASD and TD individuals using parameters consistent with those used in the newborn blood analysis **(Table 4) (Supplemental Figure 8) (Supplemental Tables 15-23)**. Similar to newborn blood, DMRs were more often hypomethylated than hypermethylated in ASD compared to TD samples in females, while in males, this finding was inconsistent across tissues **(Supplemental Figure 9)**. While there was significant overlap between DMRs identified between females and males in each tissue, loci were often methylated in opposite directions in the two sexes **(Supplemental Table 24)**.

**Table 4.**
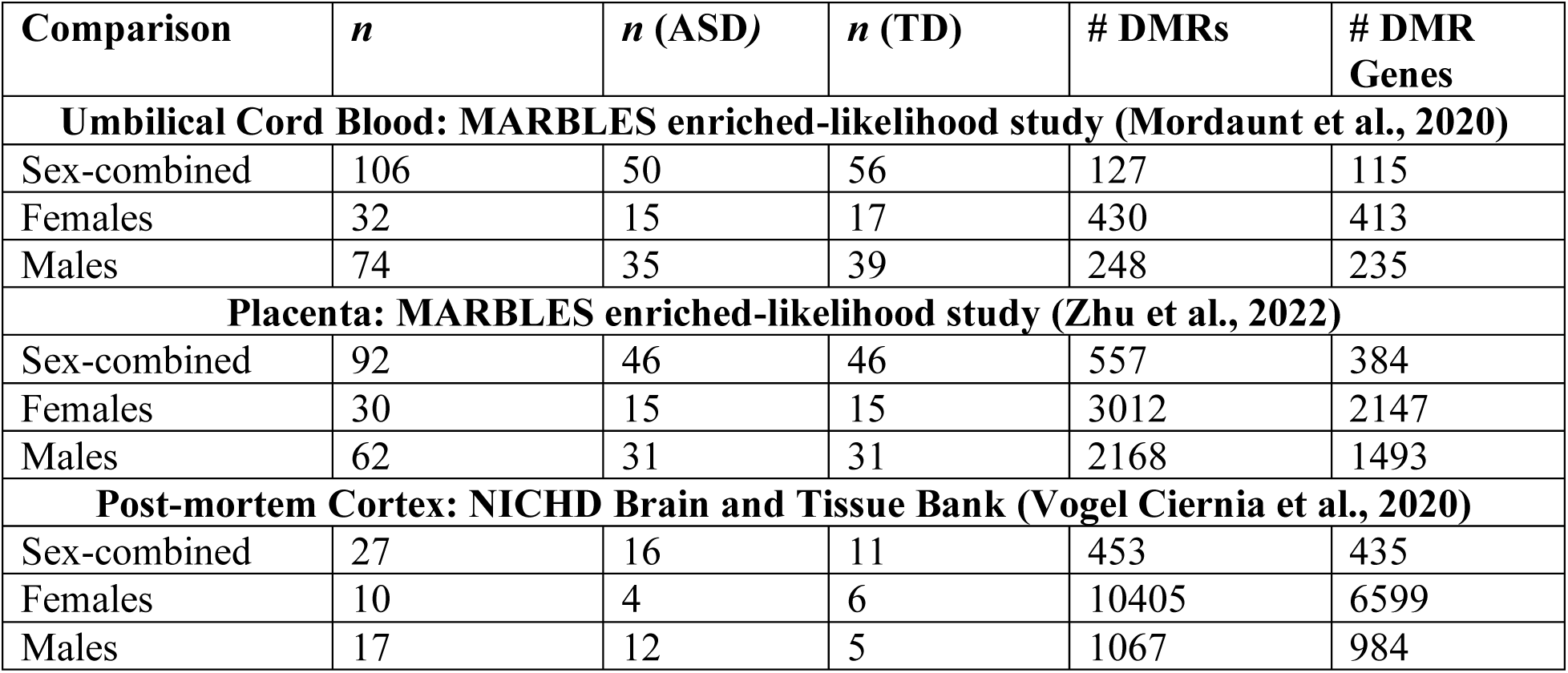
ASD vs TD DMRs in cord blood, placenta, and cortex.

| Comparison | *n* | *n* (ASD) | *n* (TD) | # DMRs | # DMR Genes |
| --- | --- | --- | --- | --- | --- |
| **Umbilical Cord Blood: MARBLES enriched-likelihood study (Mordaunt et al., 2020)** |  |  |  |  |  |
| Sex-combined | 106 | 50 | 56 | 127 | 115 |
| Females | 32 | 15 | 17 | 430 | 413 |
| Males | 74 | 35 | 39 | 248 | 235 |
| **Placenta: MARBLES enriched-likelihood study (Zhu et al., 2022)** |  |  |  |  |  |
| Sex-combined | 92 | 46 | 46 | 557 | 384 |
| Females | 30 | 15 | 15 | 3012 | 2147 |
| Males | 62 | 31 | 31 | 2168 | 1493 |
| **Post-mortem Cortex: NICHD Brain and Tissue Bank (Vogel Ciernia et al., 2020)** |  |  |  |  |  |
| Sex-combined | 27 | 16 | 11 | 453 | 435 |
| Females | 10 | 4 | 6 | 10405 | 6599 |
| Males | 17 | 12 | 5 | 1067 | 984 |

We mapped DMRs from all analyses to the nearest gene and overlapped gene lists across tissues, finding that discovery newborn blood DMR genes overlapped more than would be expected by chance with those in other tissues. This included umbilical cord blood in females, males, and sex-combined analyses, as well as placenta in females and males, but not cortex in any comparison **(Figure 4A, Supplemental Table 25)**. In the sex-combined comparison, no DMR genes were identified in all four tissues. In females-only, however, three genes mapped to DMRs identified in all four tissues: *BCOR* (chrX), *GALNT9* (chr12), and *OPCML* (chr11) **(Figure 4C, Supplemental Figure 10).** *BCOR* and *OPCML* were also replicated in newborn blood from females **(Supplemental Table 10)**. In males, one gene mapped to DMRs in all tissues: *ZNF733P* (chr7) **(Figure 4B)** but was not replicated in the newborn blood replication dataset.

**Figure 4.**
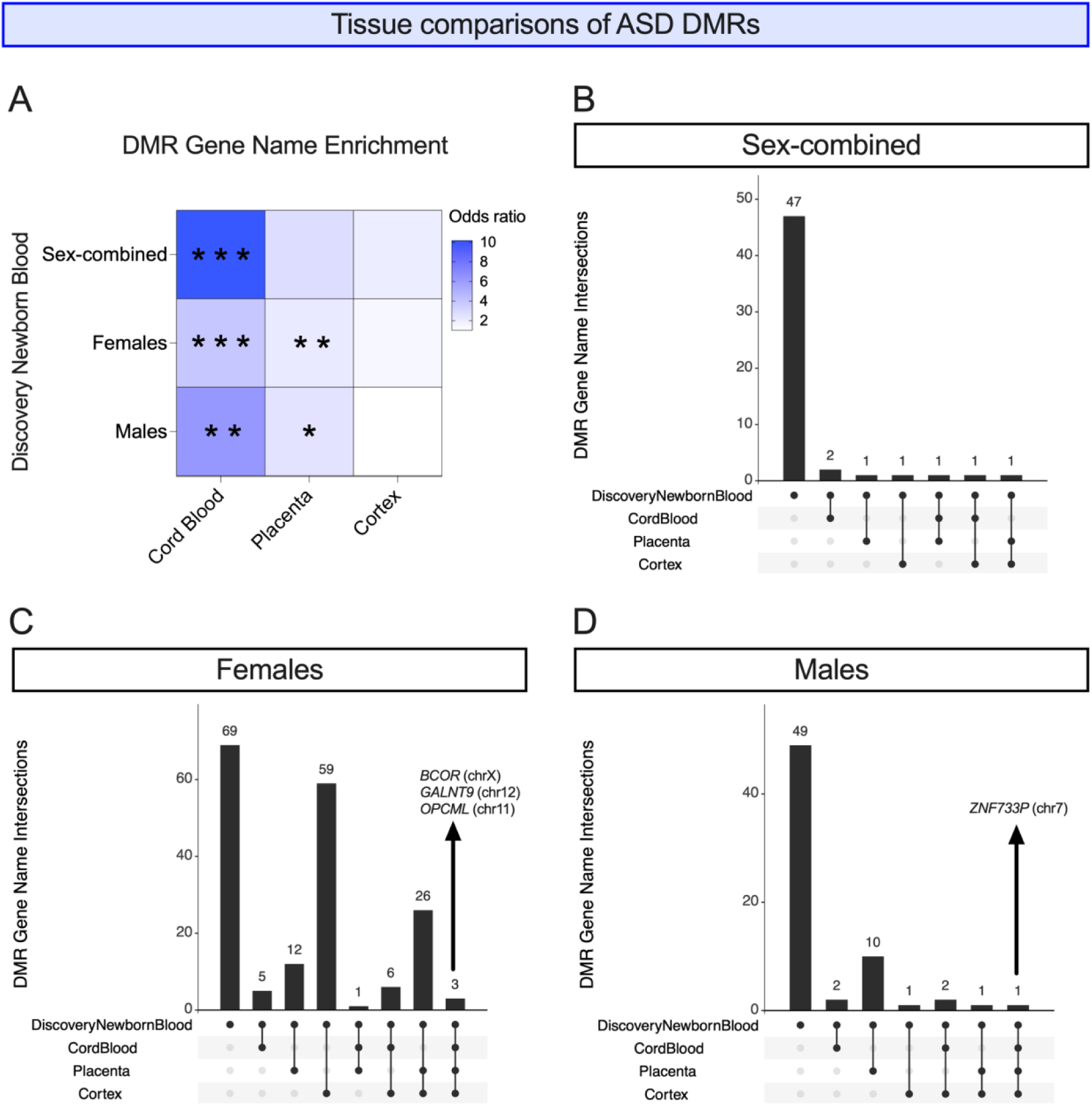
Tissue comparisons of ASD DMRs. **A)** ASD DMR genes from discovery newborn blood were overlapped with those from cord blood, placenta, and cortex from sex-combined, females only, and males only comparisons. Heatmap shows odds ratios of overlaps where darker blue indicates a higher odds ratio. * *p* <0.05, ** *p* <0.01, *** *p* <0.001, **** *p* <0.0001. UpSet Plots showing numbers of intersections between ASD DMR genes from discovery newborn blood and those from cord blood, placenta, and cortex in **B)** sex-combined, **C)** females only, and **D)** males only comparisons.

### ASD DMRs in different tissues are closer together than expected by chance, particularly in females, but are still tissue-specific

As a more stringent comparison than gene name overlap, we overlapped DMRs from newborn blood with other tissues by genomic location. Since DMRs cover a very small fraction of the genome, it is not surprising that there were zero loci that had a DMR in all four tissues in either females or males. However, in females, there were two DMRs that were detected in newborn blood and two other tissues: chr2:130037414-130037709 (chr2q21.1 band) in newborn blood (discovery), cord blood, and placenta; and chr20:30899557-30900219 (chr20q11.21 band) in newborn blood (discovery and replication), cord blood, and cortex **(Supplemental Table 26)**. We calculated the significance of tissue-tissue pairwise overlaps by genomic location and found that, in females, the overlap of newborn blood DMRs was significantly higher than expected by chance versus cord blood (*p*=9.99E-5) and placenta (*p*=0.048) **(Supplemental Table 27)**. In males, no overlaps between any two tissues were significant.

As the likelihood of DMRs overlapping by genomic locations across analyses is very low, we also tested whether DMRs identified in different tissues were closer together on the genome than would be expected by chance. We found that for all pairwise tissue comparisons of DMRs identified from females, the mean distance of a DMR in newborn blood was closer to the nearest DMR in all other tissues than would be expected by chance (all permutation *p*-values <0.001) **(Supplemental Figure 11) (Supplemental Table 27)**. In males, ASD DMRs from newborn blood were closer by chance with DMRs from cord blood (permutation *p*-value <0.001), but farther than expected from DMRs in cortex (permutation *p*-value =0.0001).

Despite ASD DMRs being closer together than expected by chance across many tissues, particularly in females, we hypothesized that newborn blood DMRs are largely tissue specific and would not differentiate ASD from TD samples from other tissues. To test this tissue specificity, we assayed the smoothed CpG methylation values in discovery newborn blood samples over the DMRs identified in cord blood, placenta, and cortex and confirmed by PCA that the samples did not separate by ASD diagnosis in either sex using DMRs from any tissue **(Supplemental Figure 12).**

### ASD DMRs identified from females are enriched for neurodevelopment-related biological processes in all tissues

We next performed gene ontology **(GO)** analysis on ASD DMRs from newborn blood as well as from cord blood, placenta, and cortex to evaluate whether consistent processes were epigenetically dysregulated across tissues. We selected all biological processes with *p*<0.2 from each DMR comparison for overlap and found 15 terms that appeared in all tissues’ female DMR analyses, with the most significant term (on average across tissues) being “central nervous system neuron differentiation” **(Figure 5A)**. Other terms identified in females relevant to neurodevelopment or epigenetics included “ventral spinal cord interneuron specification” and “regulation of histone H3-K36 methylation”. In males, only six terms appeared in all tissues and were related to GTPase activity and muscle cell migration.

**Figure 5.**
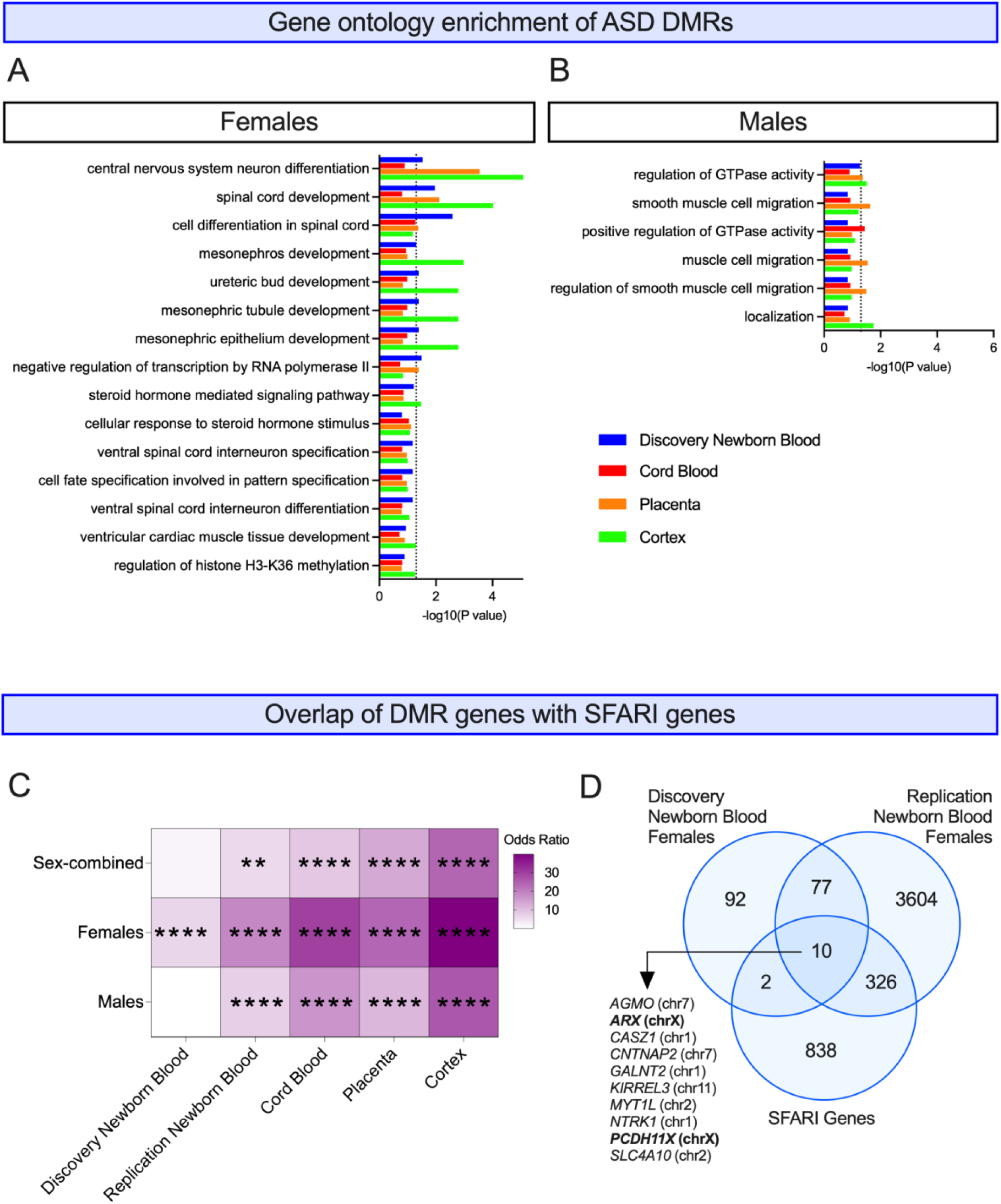
Enrichment of ASD DMRs with biological processes and ASD-risk genes. Gene ontology enrichment in **A)** females and **B)** males. Sex-specific DMRs from each tissue were examined for gene ontology enrichments using GOfuncR. Biological process enrichments with *p* < 0.2 from all tissues were overlapped and graphed by their -log(p-value). The dotted black lines represent *p* = 0.05. **C)** Heatmap of sex-specific DMRs from each tissue overlapped with SFARI genes, with darker purple indicating a higher odds ratio (range: 0-40). * *p* <0.05, ** *p* <0.01, *** *p* <0.001, **** *p* <0.0001. **D)** Venn diagram of DMR genes identified in females from discovery newborn blood and replication newborn blood overlapped with SFARI genes.

### ASD DMR genes significantly overlap with SFARI genes, particularly in females

Because DMRs from all tissues were enriched for neurodevelopmental processes, we hypothesized that they were also enriched for known ASD-associated genes. To test this, we overlapped DMRs mapped to genes with all genes in the SFARI database (46). In newborn blood, only females were significantly enriched (*p*<0.05) for SFARI genes, while in all other tissues, females, males, and sex-combined DMRs were significantly enriched for SFARI genes **(Figure 5B, Supplemental Table 28)**.

To identify replicated newborn blood loci that have been genetically linked with ASD, we overlapped discovery and replication newborn blood DMR genes with SFARI genes. We found that of the 87 newborn blood DMR genes that replicated in females across two cohorts, 10 (11.5%) were listed as SFARI genes, two of which are X-linked **(Figure 5C)**: *AGMO* (chr7), *ARX* (chrX), *CASZ1* (chr1), *CNTNAP2* (chr7), *GALNT2* (chr1), *KIRREL3* (chr11), *MYT1L* (chr2), *NTRK1* (chr1), *PCDH11X* (chrX), and *SLC4A10* (chr2). No replicated DMRs genes identified from males were listed as SFARI genes.

## Discussion

In this study, we performed the first sex-stratified epigenomic signature of ASD from newborn blood spots, finding significant replication with an independent cohort, support for the female protective effect, and evidence of biological relevance of the gene loci and gene pathways identified. Our findings will help inform future efforts to develop newborn screening tools for ASD, improve our understanding of the etiology and pathology of ASD, and highlight the importance of sex-stratification in ASD studies.

Our ASD DMRs from discovery newborn blood revealed a signature of hypomethylation in ASD compared to TD samples as well as enrichment for DMR location in promoters, CpG islands, and CpG shores in both sexes, findings that have been previously reported in cord blood (17). Despite the similarities in epigenetic profiles across the sexes, only one DMR gene was identified in all three comparisons (sex-combined, females only, and males only): *SPATA19*, which has been previously implicated in dysregulated neurology and behavior (47,48), but not in ASD specifically. These findings indicate that, despite similar patterns of ASD signatures, the dysregulated loci themselves may be different in females and males, demonstrating the value in sex-stratifying samples for ASD studies, despite reduced sample sizes.

Our analysis of sex differences in ASD provides the first epigenetic support for the female protective effect, which hypothesizes that the protective nature of certain factors in females (such as DNA methylation marks) require more changes to be attenuated to result in ASD, leading to fewer cases in females and stronger biological signatures. Our findings supported this model, given that we found a stronger epigenetic signature of ASD in females over regions whose methylation levels differ by sex. We also found a striking overlap between female vs male DMRs in ASD individuals and TD individuals, adding to the evidence that the mechanisms governing sex differentiation in neurotypical brains may also contribute to sex differences in ASD. We also that these biological differences are not only measurable in the brain but are reflected by methylation changes in newborn blood. The female protective effect is supported by neuroimaging studies, in which comparisons of ASD vs TD individuals have shown larger differences in females than in males with regard to cortical thickness and development (49), organization of nerve fibers (50,51), gray matter asymmetry (52), amygdala functional connectivity with the cortex (53), local connectivity within brain networks (54). Similarly, genetic studies that have found that females with ASD carry more autism-associated genetic mutations than do males with ASD (55–59). To our knowledge, this is the first DNA methylation study to examine proposed models for sex differences in ASD and report epigenetic support for the female protective effect.

These female-driven sex differences in ASD that we observed from autosomal DMRs may be augmented by the enrichment of female-specific ASD vs TD DMRs on the X chromosome, which we observed in both cohorts of newborn blood. Interestingly, enrichment for ASD DMRs on the X chromosome has previously been reported in umbilical cord blood in both males and females (17), while our finding was restricted to females. Rather than reflecting a tissue difference, this finding may be due to differences in genetic risk for ASD (which is reflected in epigenetics) in the different cohort types. The discovery and replication newborn blood datasets are from the population-based CHDS birth cohort and the CHARGE case-control study, while the cord blood was obtained from the MARBLES enriched-likelihood cohort, which enrolled pregnant women who already had a child with ASD (25). Therefore, the MARBLES mothers may be protected carriers of genetic and/or epigenetic pre-disposition for ASD, including on the X chromosome, which then would likely have been reflected in enrichment for X-linked DMRs in both sexes.

Having established the sex specificity of the epigenetic signature of ASD, we hypothesized that sex-specific DNA methylation differences may be detectable across multiple perinatal tissues if the dysregulation occurred prior to tissue differentiation, lending insight to etiology and pathogenesis. In females, one or more DMRs from all four tissues mapped to *BCOR*, *GALNT9*, and *OPCML*, all of which have linked to neuropathology or dysregulated neurodevelopment (60–67). It is striking that all three genes that replicated across tissues in females are linked to neurodevelopment generally or ASD specifically, given that three out of four tissues are collected at the time of birth and are not directly linked to disease pathogenesis. *ZNF733P*, identified from all tissues in males, is a pseudogene located nearby many zinc-finger protein-encoding genes close to the centromere of chromosome 7 and to our knowledge, has not previously been associated with neurodevelopmental disorders.

We also found evidence for the biological relevance of the epigenetic signature of ASD in newborn blood and other perinatal tissues. Females in particular showed enrichment of DMRs from newborn blood, cord blood, placenta, and cortex for neurodevelopment-related biological processes, such as “central nervous system neuron differentiation”. In addition, ASD DMRs across all tissues and sexes were enriched for SFARI genes, indicating convergence of genetic and epigenetic markers of ASD and aligning with the findings from a 2021 study, in which autism-associated DNA CpG methylation sites from maternal blood, cord blood, and placenta samples were enriched for SFARI genes (20). Overall, these results show the biological relevance of perinatal tissues as surrogates for the brain in understanding epigenetic dysregulation in ASD.

In addition to furthering the understanding of sex-specific epigenetic dysregulation in ASD, our results provide insights for biomarker development. Reproducible DNA methylation differences detectable in newborn blood could be potentially added to molecular screening panels to identify infants who should be behaviorally screened for ASD. Newborn biomarkers may be particularly useful for cases that are harder to identify behaviorally, and may help to overcome sex biases and other disparities in ASD diagnosis (68). A positive screening result would inform parents and medical practitioners that diagnostic services will be paramount for the child and, importantly, a negative screening result would not exclude the possibility of future diagnosis. The possibility of using newborn blood for screening is particularly plausible given the accessibility and widespread collection of NDBS. While additional research is needed in this area, our study provides initial results supporting a possible future of ASD screening in newborns using DNA methylation from newborn blood.

### Limitations

Despite its strength as the first WGBS study of ASD in newborn blood, this study was limited by its sample sizes, due the inherent challenge in obtaining a sufficient number of perinatal samples from individuals later diagnosed with ASD. Consistent with most ASD studies, our samples have a strong male bias, which may influence the sex-specific findings. When we down-sampled males to match the number of females in discovery newborn blood, the number of DMRs were comparable across sexes, indicating that the lower number of male DMRs across tissues was an artifact of higher sample sizes, despite our adjustment of the “perGroup” variable to try to account for this. This may have affected downstream analyses, such as overlap with SFARI genes, where we found significant enrichment across all tissues in both sexes except in discovery newborn blood, where sex-combined and males-only comparisons did not show enrichment, potentially because there were fewer DMRs in these comparisons than for females or other tissues. Another potential limitation arises from our threshold of permutation *p*-value<0.05 for DMR identification. We did not correct DMR *p*-values for multiple hypotheses because there is high correlation of methylation across regions, meaning that each DMR does not represent a single hypothesis (69). This purposely lenient threshold reduced the chance of false negatives, though we acknowledge that this strategy also likely increased the number of false positive results. We addressed this limitation by using a replication group for newborn blood, and by comparing the DMRs across multiple tissues. Lastly, there may be cell type heterogeneity amongst our samples that influenced the DNA methylation signatures. We did not adjust DMRs for cell types because estimations from different tools and reference datasets have shown large inconsistencies when estimating cell type proportions from WGBS data, particularly from NDBS, thereby decreasing our confidence in using cell type as an adjustment variable (33). Additionally, a previous ASD cord blood study showed that when nRBCs were adjusted for, a significant majority of ASD DMRs were maintained (17). Instead, we used a comparison of newborn blood to four other tissues to identify pan-tissue ASD DMRs that would be unlikely to be impacted by subtle cell type differences between ASD vs TD samples.

## Conclusions

This study is the first to present sex-stratified WGBS DNA methylation signatures of ASD in newborn blood. We found a signature of hypomethylation in ASD compared to TD samples in both sexes, and enrichment for DMRs in promoters, CpG islands and CpG shores. Because there was little overlap between DMR genes identified from females and males, we further explored sex differences in newborn blood and are the first to our knowledge to report epigenetic support for the female protective effect. We further found that the newborn blood ASD DMRs overlapped with those from other perinatal tissues (umbilical cord blood and placenta) as well as post-mortem cortex, showing that despite the tissue-specificity of DNA methylation there may be some loci that are consistently dysregulated across tissues, perhaps due to disruption early in life. ASD DMRs from all tissues were enriched for neuro-related biological processes (females) and SFARI ASD genes (females and males), showing the biological relevance of the dysregulated loci and the convergence of genetic and epigenetic markers for ASD. This study presents a step forward in identifying potential biomarkers of ASD in newborns as well as providing insights to sex-specific epigenetics dysregulation in ASD.

## Acknowledgements

This work was supported by National Institutes of Health NIEHS T32 ES007059 (JSM), R01 ES029213 (JML, RJS), and P30 ES023513; UC Davis Perinatal Origins of Disparities Center Graduate Student Fellowship (JSM); UC Davis Perinatal Origins of Disparities IMPACT Center, Office of Research; Department of Defense Autism Research Program W81XWH-16-1-0254 (BAC).

We thank Christine Wu Nordahl for her thorough reading of the manuscript and her expertise on sex differences in ASD.

## Availability of Data

Code for this study is available on GitHub (https://github.com/juliamouat/NewbornBlood_DNAmethylation_ASD). The WGBS data from cord blood, placenta, and cortex are available in the Gene Expression Omnibus with accession numbers GSE140730 (cord blood), GSE178206 (placenta), GSE81541 (cortex) and GSE119981 (cortex). The WGBS data from the discovery and replication newborn blood are not publicly available: Any uploading of genomic data (including genome-wide DNA methylation data) and/or sharing of California Biobank Program biospecimens or individual data derived from these biospecimens has been determined to violate the statutory scheme of the California Health and Safety Code Sects. 124980(j), 124991(b), (g), (h), and 103850 (a) and (d), which protect the confidential nature of biospecimens and individual data derived from biospecimens. Should we be contacted regarding individual-level data contributing to the findings reported in this study, inquiries will be directed to the California Department of Public Health Institutional Review Board to establish an approved protocol to utilize the data, which cannot otherwise be shared peer-to-peer.

