## Supplemental Figures for "Sex-specific DNA methylation signatures of autism spectrum disorder in newborn blood"

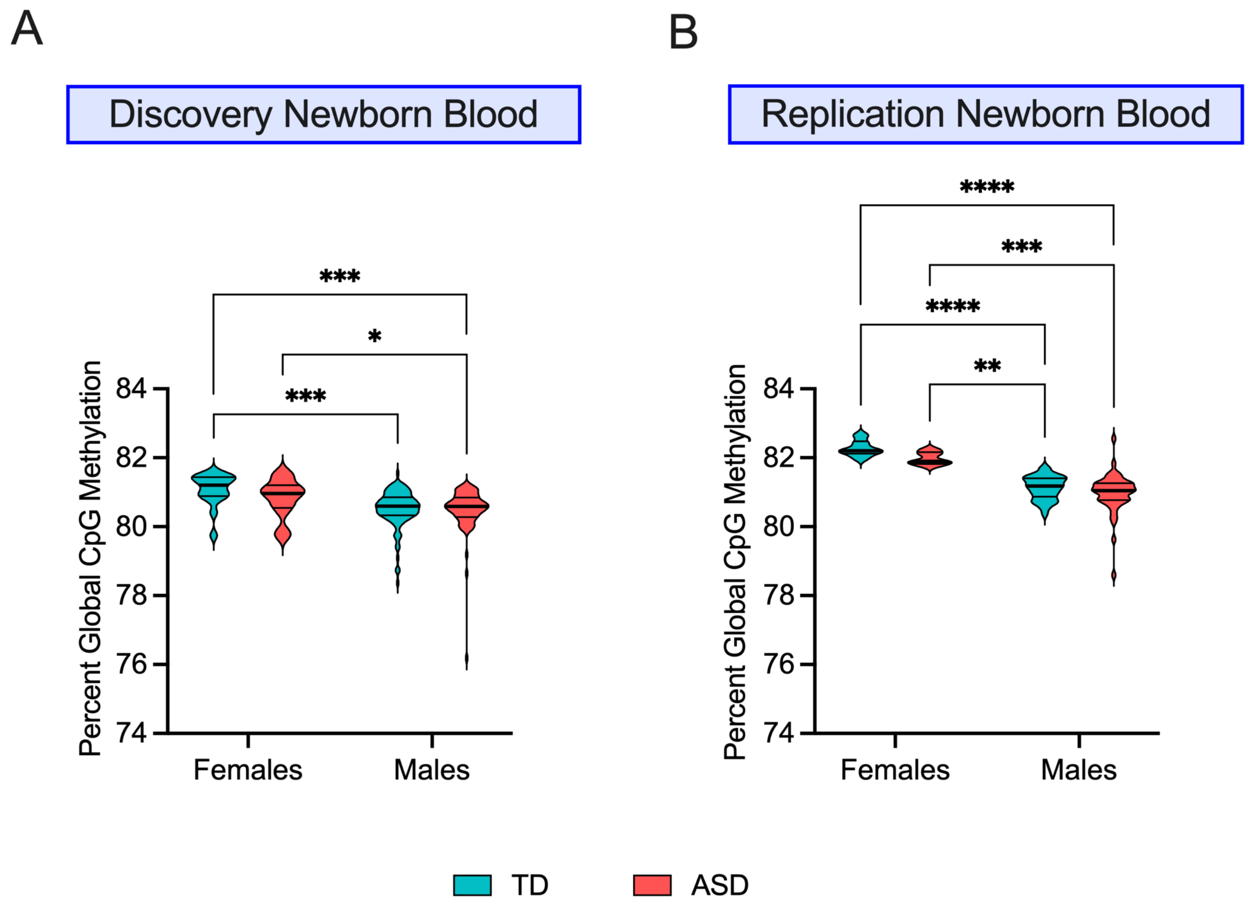


**Supplemental Figure 1.** Percent global CpG methylation in **A)** discovery and **B)** replication newborn blood, separated by sex and ASD diagnosis. Significance was tested using two-way ANOVA with Fisher's Least Significant Difference test for individual comparisons (* $p < 0.05$, ** $p < 0.01$, *** $p < 0.001$, **** $p < 0.0001$).

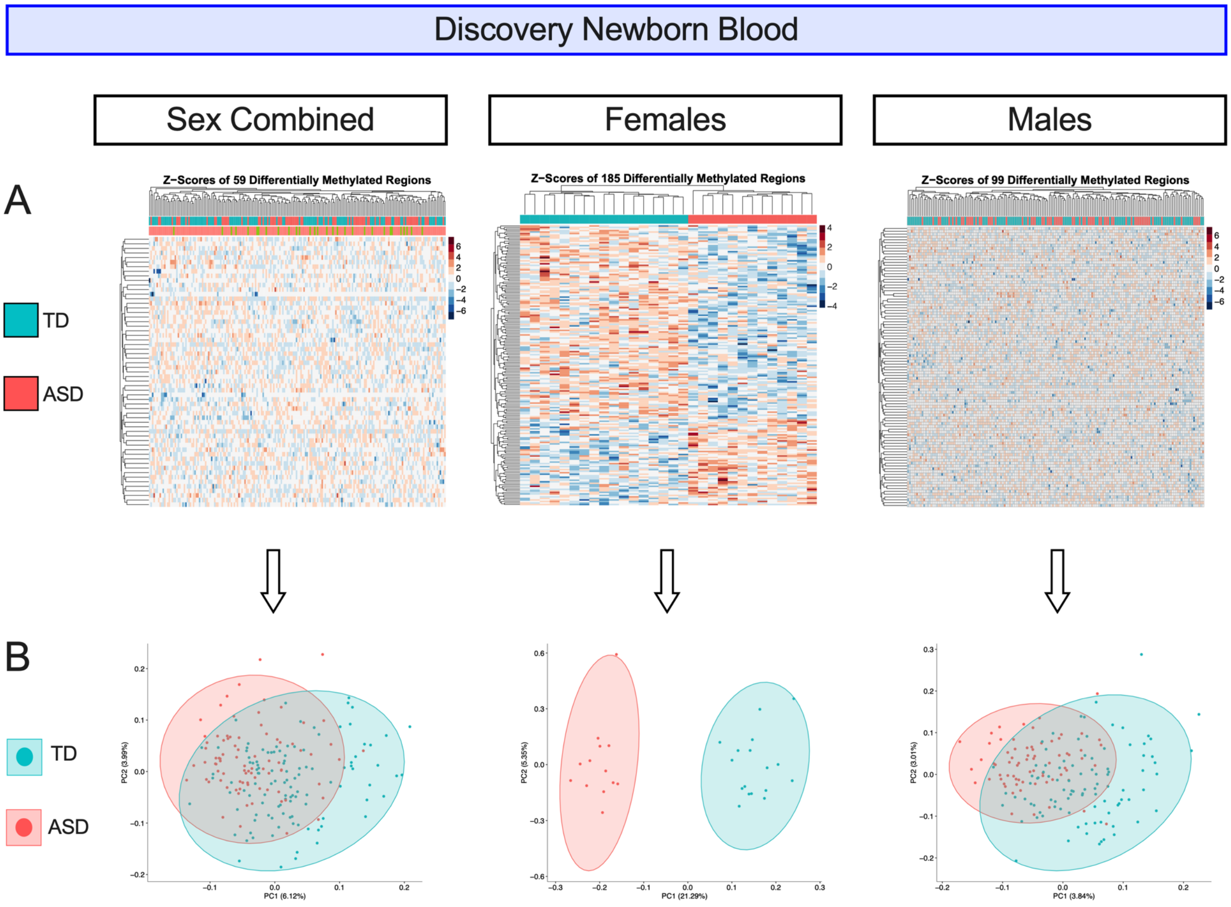


**Supplemental Figure 2.** Differentially methylated regions from sex-combined, females only, and males only comparisons of ASD vs TD samples in discovery newborn blood. **A)** Heatmaps of nominally significant ($p < 0.05$) DMRs, which are hierarchically clustered by their Z-scores, the number of standard deviations from the mean smoothed methylation value for each DMR. **B)** PCA plots of smoothed methylation values of the DMRs, colored by ASD diagnosis. The semi-transparent circles on the PCA plots represent 95% confidence intervals.

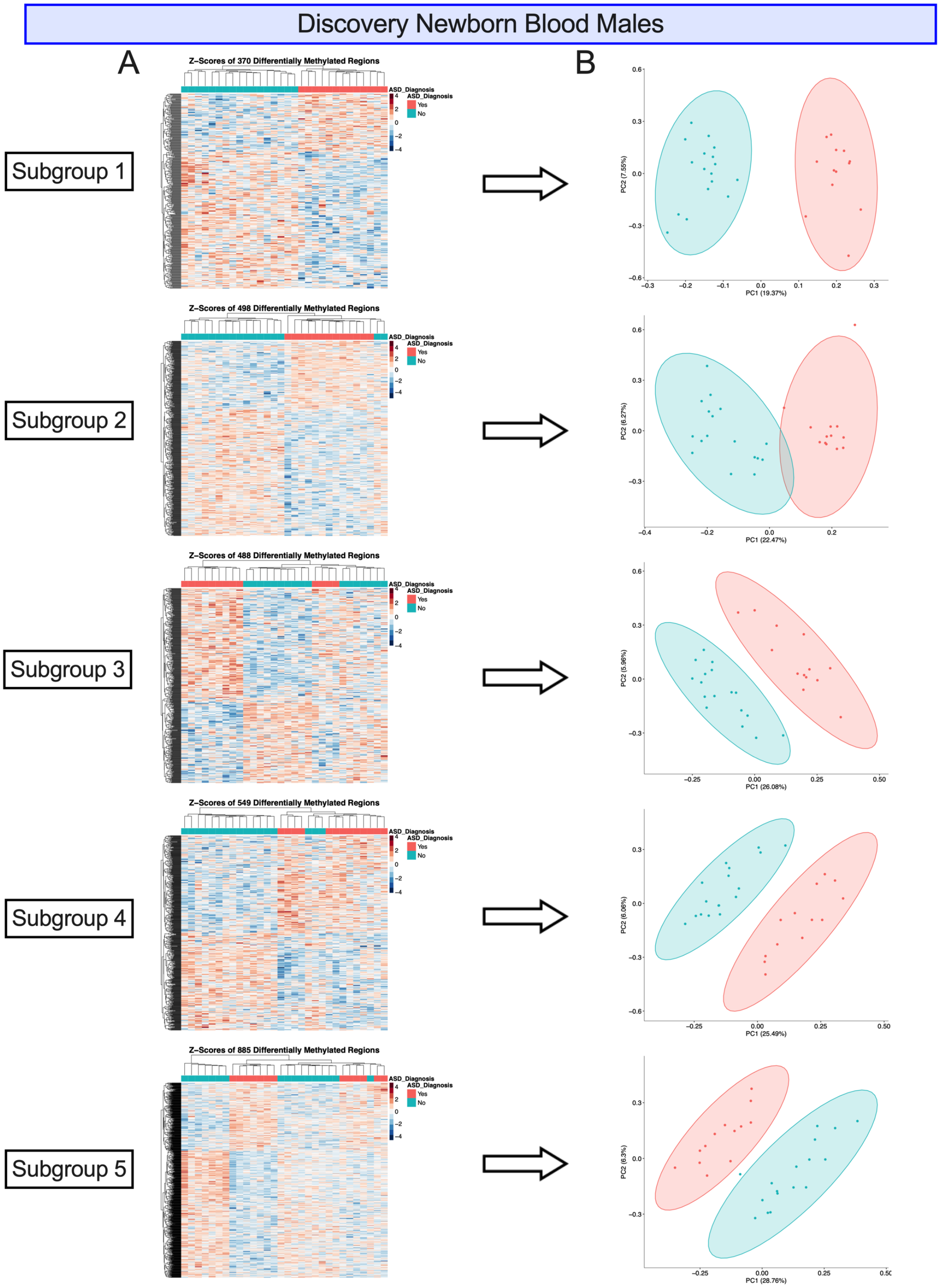


**Supplemental Figure 3.** Differentially methylated regions from discovery newborn blood ASD vs TD males. Each subgroup contains 30 randomly selected, non-repeated samples (17 TD, 13 ASD) to match the number of discovery newborn blood females. **A)** Heatmaps of nominally significant ($p < 0.05$) DMRs, which are hierarchically clustered by their Z-scores, the number of standard deviations from the mean smoothed methylation value for each DMR. **B)** PCA plots of smoothed methylation values of the DMRs, colored by ASD diagnosis. The semi-transparent circles on the PCA plots represent 95% confidence intervals.

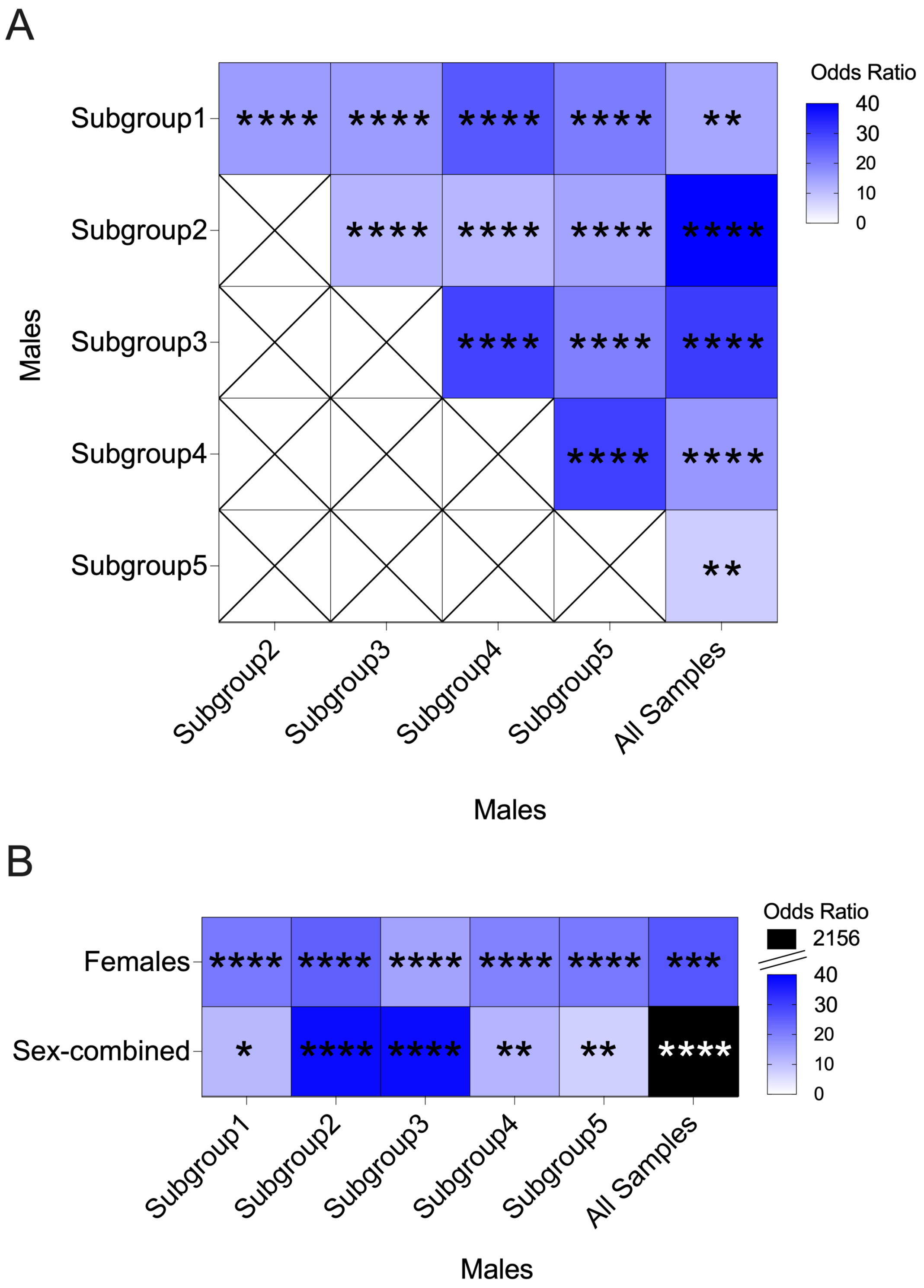


**Supplemental Figure 4.** Odds ratios of overlaps of discovery newborn blood ASD DMR gene names from the five subgroups each containing 30 males with **A)** one another and the ASD DMRs from all males and **B)** with ASD DMRs from the females only and sex-combined comparisons. Heatmap shows odds ratios of overlaps where darker blue indicates a higher odds ratio. * $p < 0.05$, ** $p < 0.01$, *** $p < 0.001$, **** $p < 0.0001$.

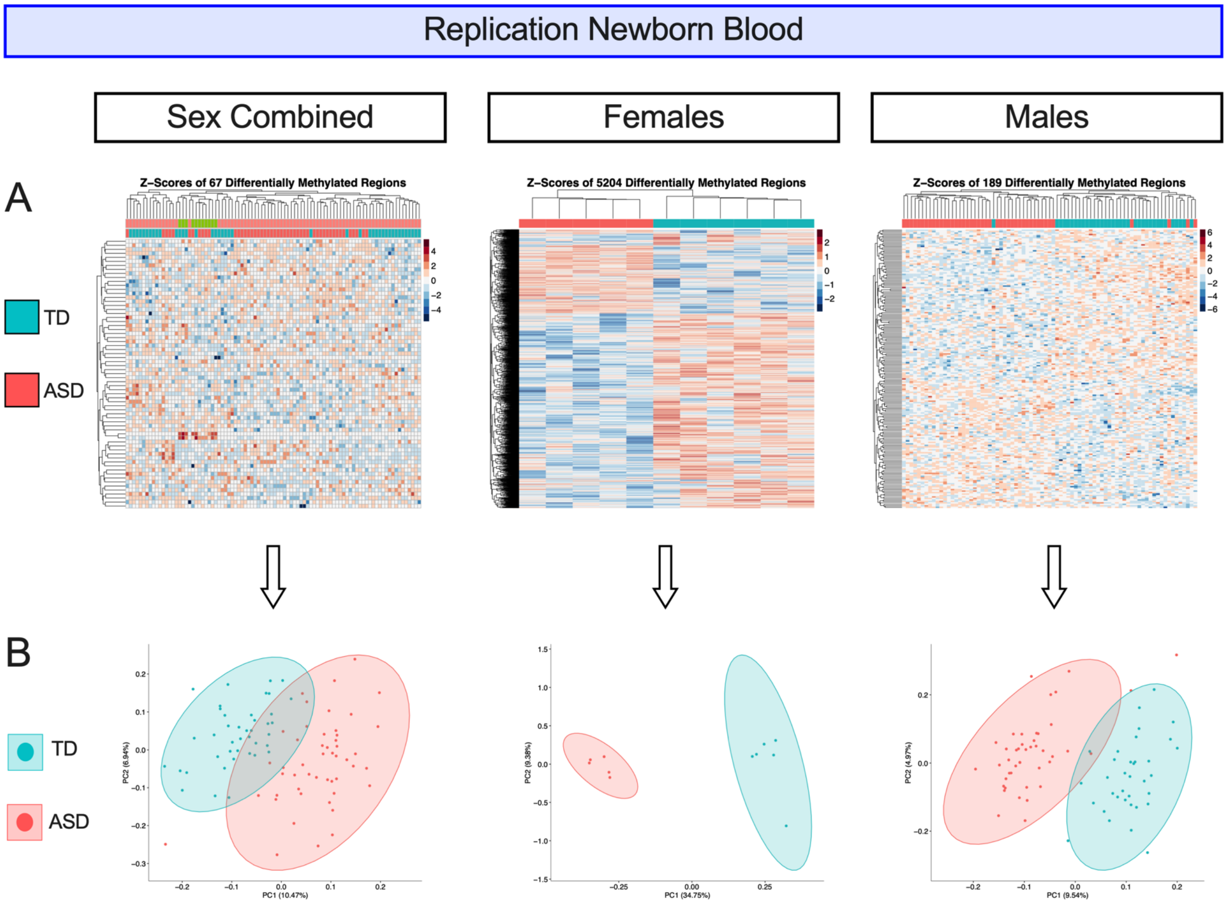


**Supplemental Figure 5.** Differentially methylated regions from sex-combined, females only, and males only comparisons of ASD vs TD samples in replication newborn blood. **A)** Heatmaps of nominally significant ($p < 0.05$) DMRs, which are hierarchically clustered by their Z-scores, the number of standard deviations from the mean smoothed methylation value for each DMR. **B)** PCA plots of smoothed methylation values of the DMRs, colored by ASD diagnosis. The semi-transparent circles on the PCA plots represent 95% confidence intervals.

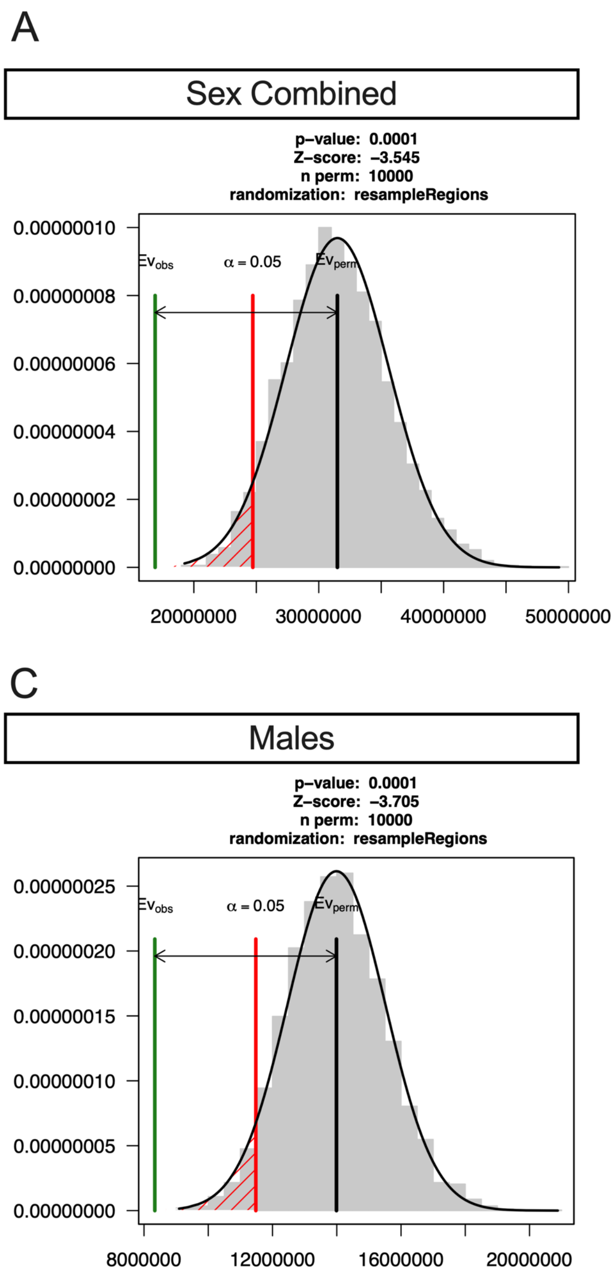


**Supplemental Figure 6.** Pairwise comparisons of the proximity of any given ASD DMR in discovery newborn blood to the closest ASD DMR in replication newborn blood in **A)** sex-combined, **B)** females only and **C)** males only comparisons using 10,000 permutations of randomly resampling regions. The black line is the expected distance, the red line is alpha = 0.05, and the green line is the observed distance.

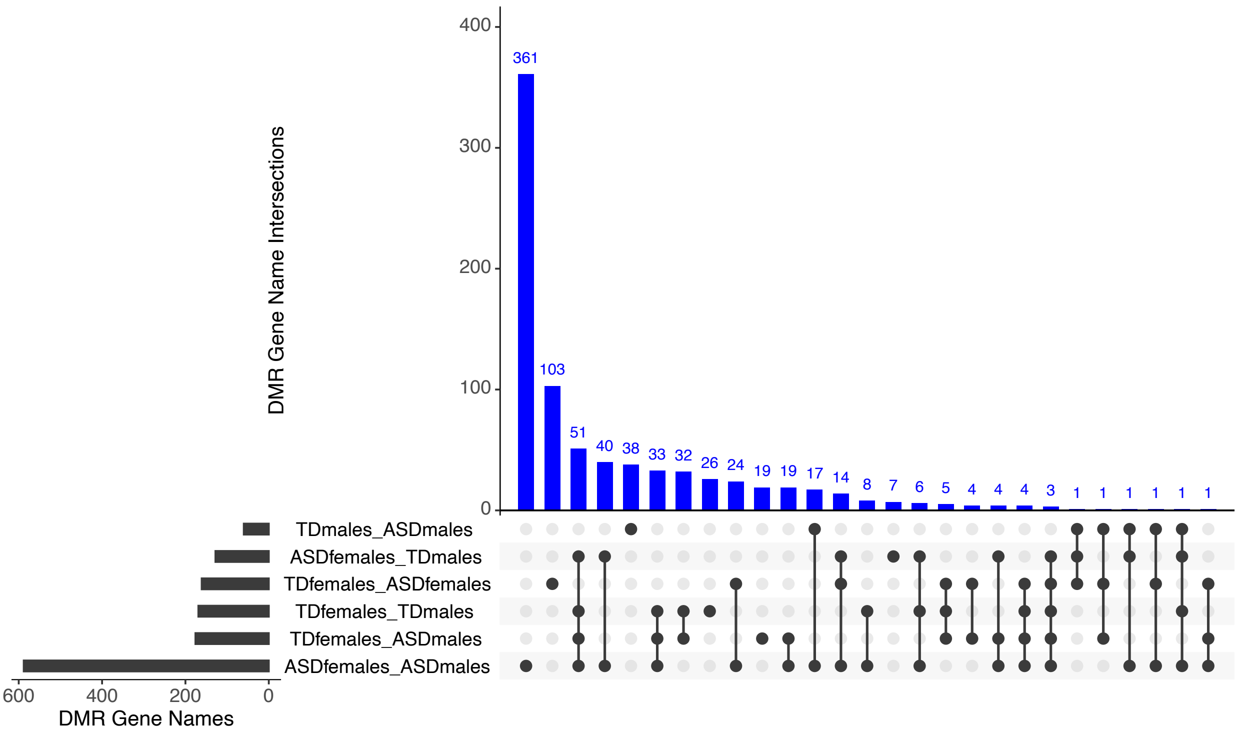


**Supplemental Figure 7.** UpSet Plot of autosomal DMRs in discovery newborn blood intersected by gene name.

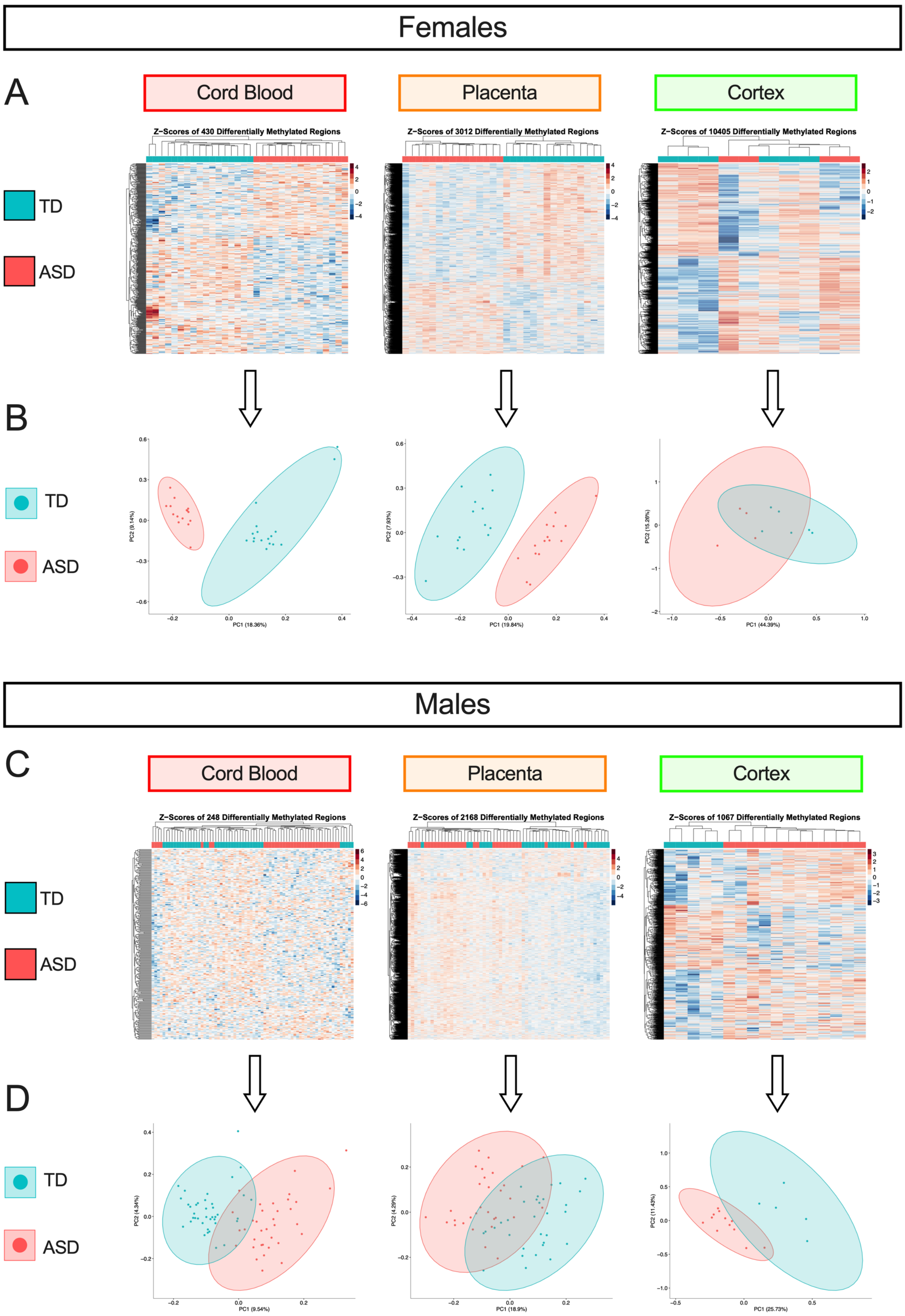

Females
A
Cord Blood
Placenta
Cortex
Z-Scores of 430 Differentially Methylated Regions
Z-Scores of 3012 Differentially Methylated Regions
Z-Scores of 10405 Differentially Methylated Regions
TD
ASD
B
TD
ASD
PC1 (18.36%)
PC2 (9.14%)
PC1 (19.84%)
PC2 (7.93%)
PC1 (44.39%)
PC2 (15.26%)
Males
C
Cord Blood
Placenta
Cortex
Z-Scores of 248 Differentially Methylated Regions
Z-Scores of 2168 Differentially Methylated Regions
Z-Scores of 1067 Differentially Methylated Regions
TD
ASD
D
TD
ASD
PC1 (9.54%)
PC2 (4.34%)
PC1 (18.9%)
PC2 (4.29%)
PC1 (25.73%)
PC2 (11.43%)

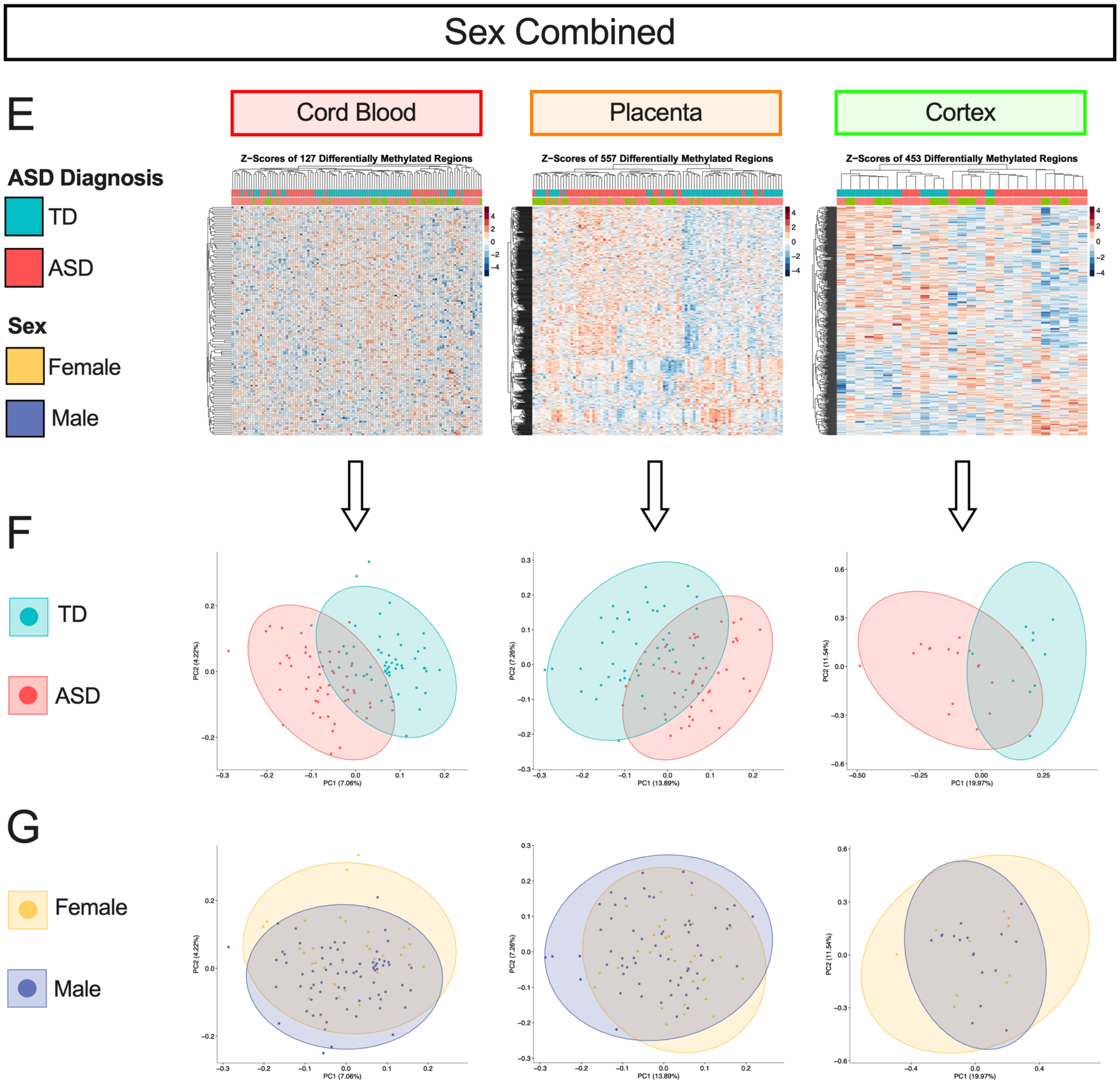


**Supplemental Figure 8.** Differentially methylated regions from ASD vs TD samples in cord blood, placenta, and cortex in females only, males only, and sex-combined (corrected for sex) comparisons. **A)** From females only, heatmaps of nominally significant (*p* <0.05) DMRs that are hierarchically clustered by their Z-scores, which are the number of standard deviations from the mean smoothed methylation value for each DMR. **B)** From females only, PCA plots of smoothed methylation values of the DMRs, colored by ASD diagnosis. The semi-transparent circles represent 95% confidence intervals .**C)** same as (A) in male samples. **D)** same as (B) in male samples. **E)** same as (A) in sex-combined samples. **F)** same as (B) in sex-combined samples. **G)** same as (B) in sex-combined samples except the PCA plots are colored by sex.

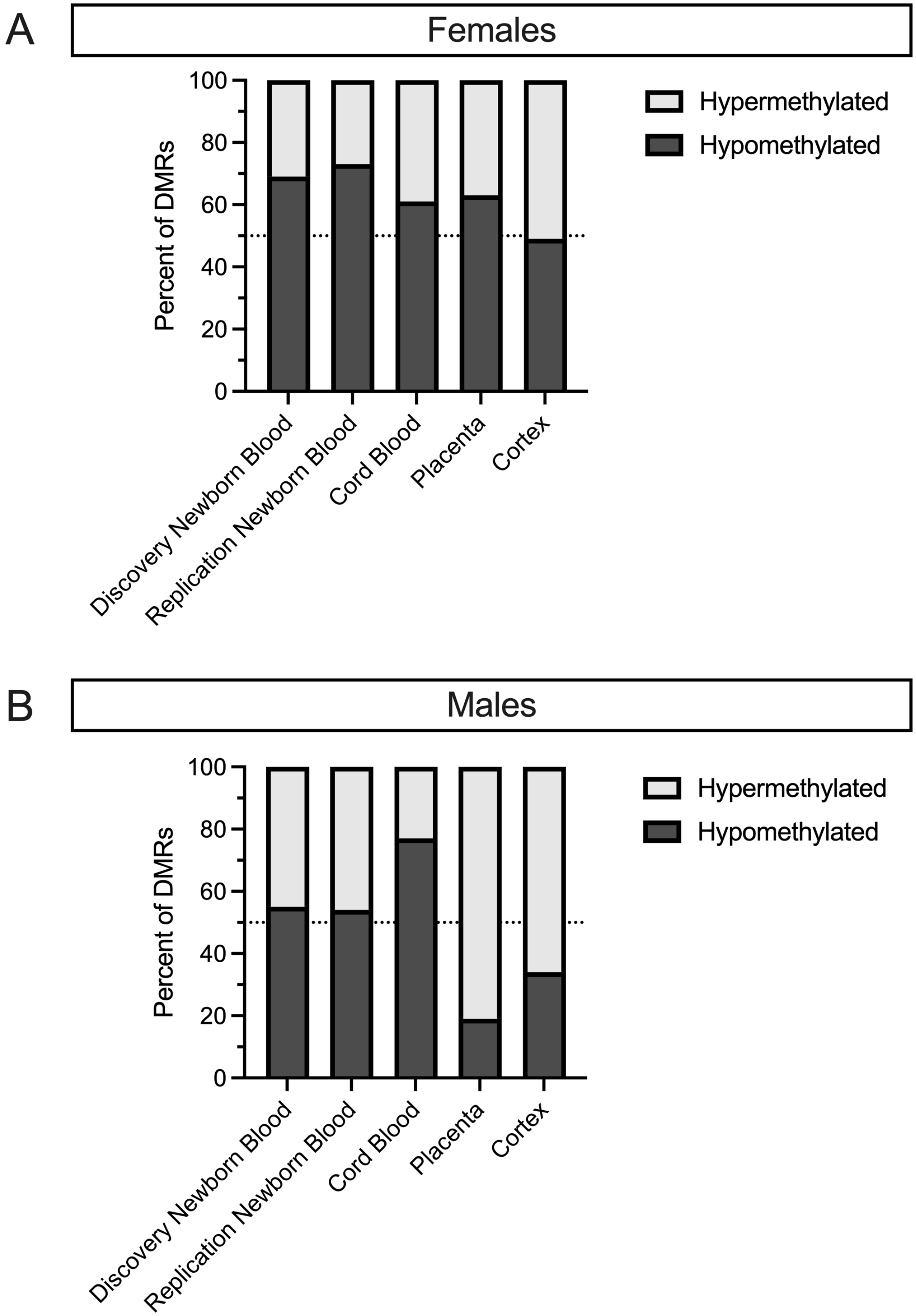


**Supplemental Figure 9.** The percent of ASD DMRs that are hypermethylated/hypomethylated in ASD vs TD samples in **A)** females and **B)** males across all tissues.

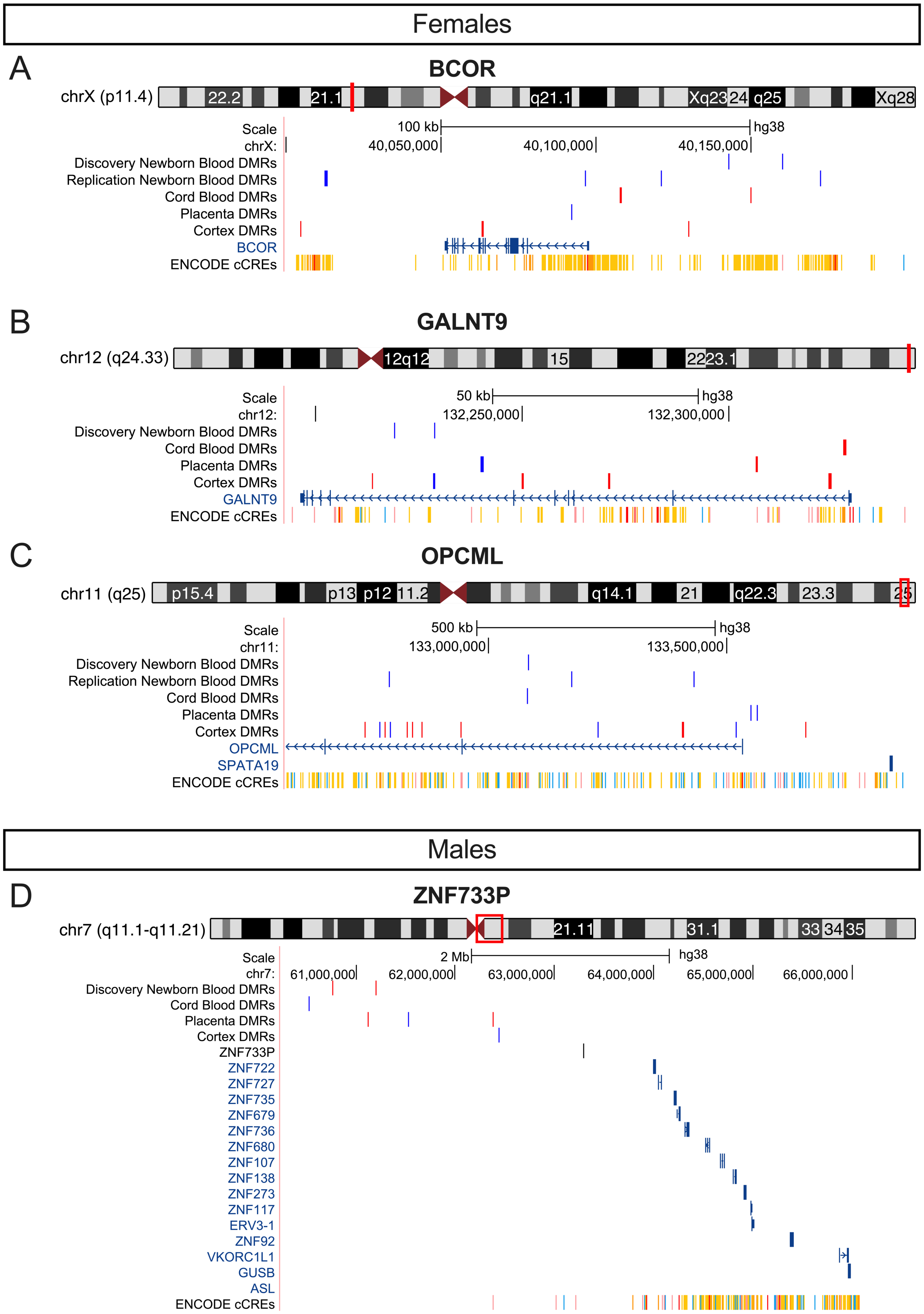


**Supplemental Figure 10. UCSC Genome Browser showing location of DMRs from all tissues** that mapped to **A)** *BCOR*, **B)** *GALNT9*, **C)** *OPCML*, and **D)** *ZNF733P*. Red lines indicate that the DMR was hypermethylated in ASD compared to TD samples with blue lines indicate hypomethylated DMRs.

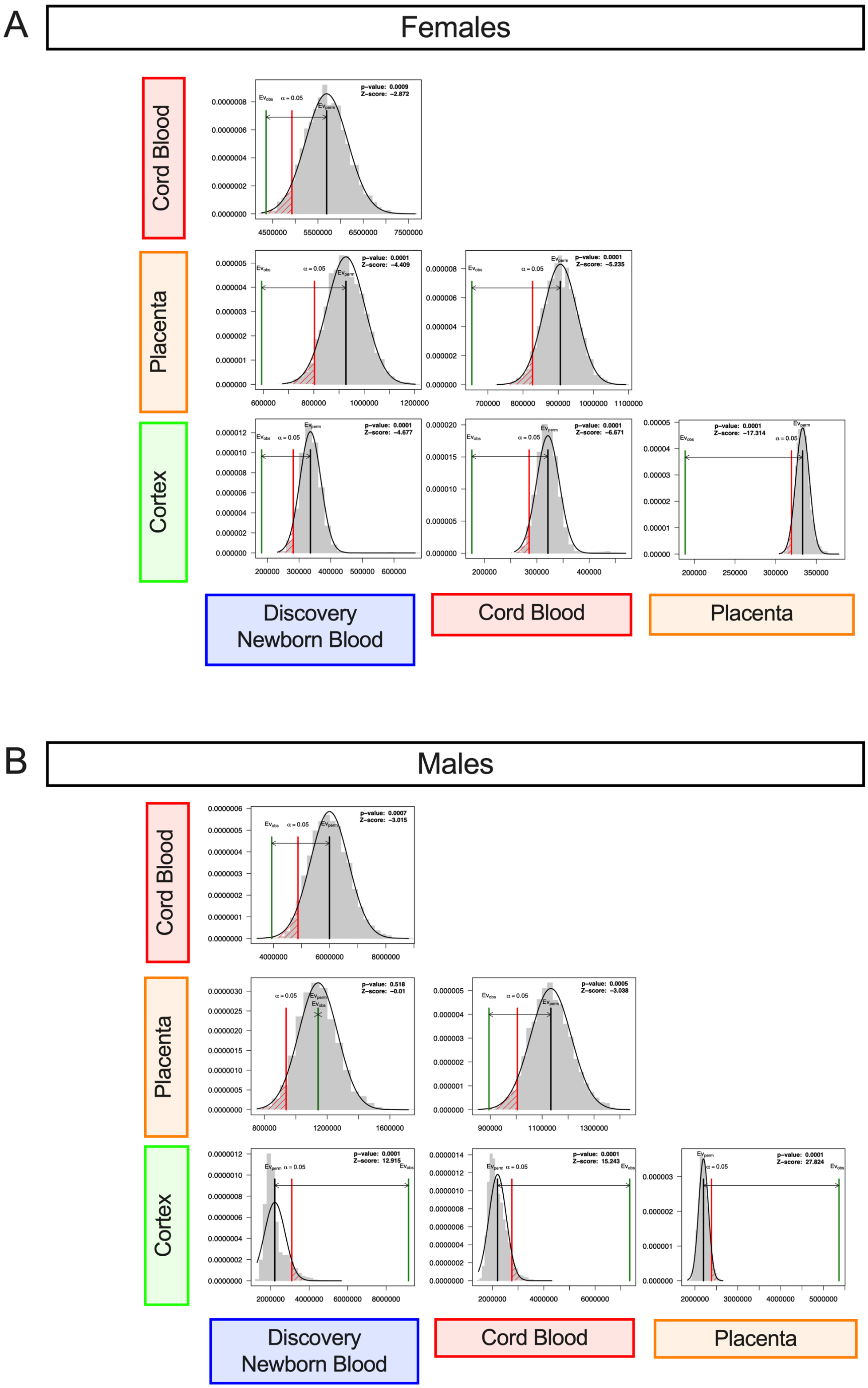


**Supplemental Figure 11.** Pairwise comparisons of the proximity of any given DMR to the closest DMR in another tissue in **A)** females and **B)** males using 10,000 permutations and random resampling of regions. The black line is the expected distance, the red line is alpha = 0.05, and the green line is the observed distance.

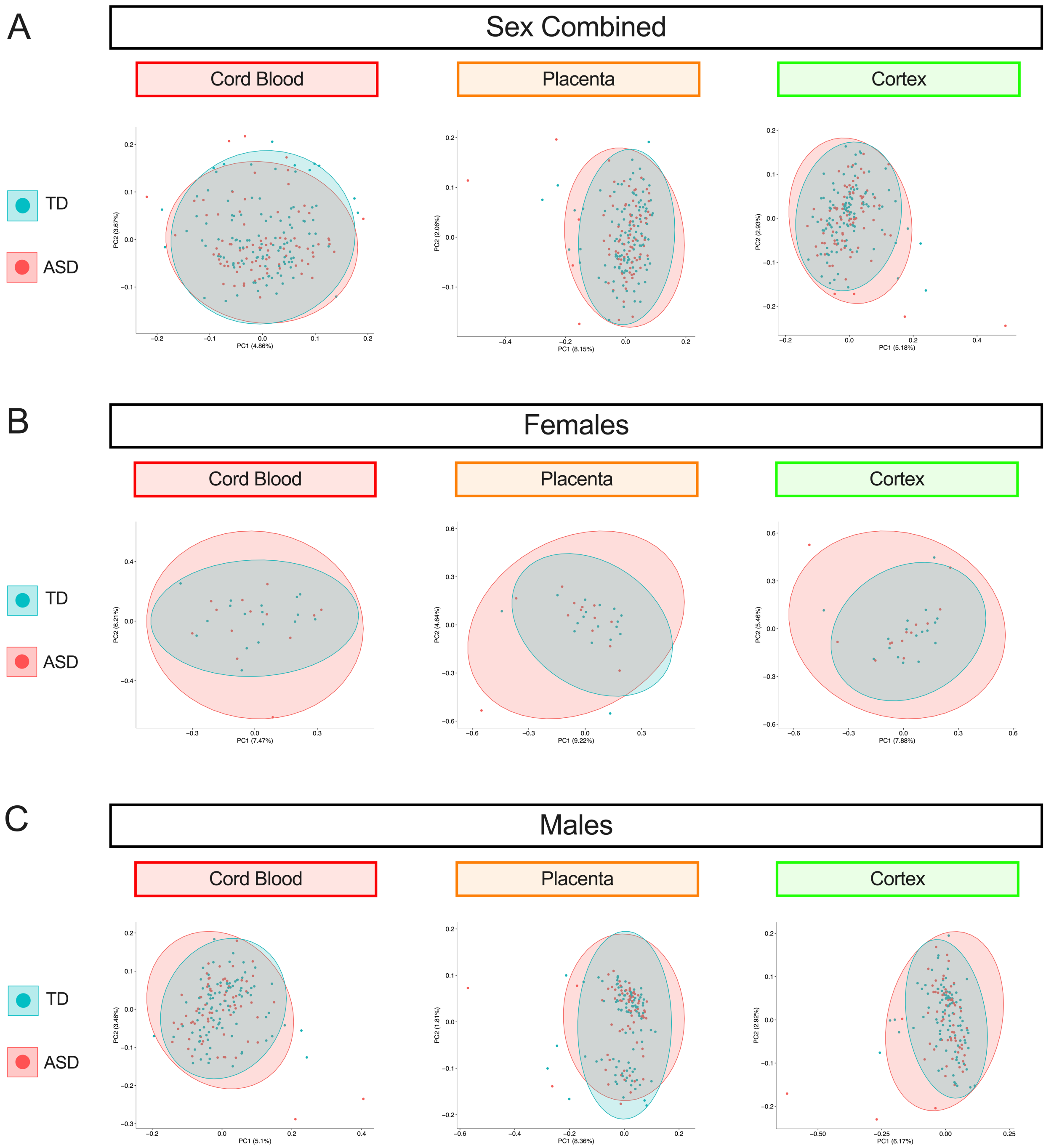


**Supplemental Figure 12.** PCA plots of smoothed methylation values in discovery newborn blood over DMRs identified from cord blood, placenta, and cortex in **A)** sex-combined, **B)** females only, and **C)** males only comparisons. The semi-transparent circles on the PCA plots represent 95% confidence intervals.
